# Dectin-1 mediated DC-SIGN Recruitment to *Candida albicans* Contact Sites

**DOI:** 10.1101/2020.12.15.422964

**Authors:** Rohan P. Choraghe, Aaron K. Neumann

## Abstract

At host-pathogen contact sites with *Candida albicans*, Dectin-1 activates pro-inflammatory signaling, while DC-SIGN promotes adhesion to the fungal surface. We observed that Dectin-1 and DC-SIGN collaborate to enhance capture/retention of *C. albicans* under fluid shear culture conditions. Therefore, we devised a cellular model system wherein we could investigate the interaction between these two receptors during the earliest stages of host-pathogen interaction. In cells expressing both receptors, DC-SIGN was quickly recruited to contact sites (103.15% increase) but Dectin-1 did not similarly accumulate. Once inside the contact site, FRAP studies revealed a strong reduction in lateral mobility of DC-SIGN (but not Dectin-1), consistent with DC-SIGN engaging in multivalent adhesive binding interactions with cell wall mannoprotein ligands. Interestingly, in the absence of Dectin-1 co-expression, DC-SIGN recruitment to the contact was much poorer—only 35.04%. These data suggested that Dectin-1 promotes the active recruitment of DC-SIGN to the contact site. We proposed that Dectin-1 signaling activates the RHOA pathway, leading to actomyosin contractility that promotes DC-SIGN recruitment, perhaps via the formation of a centripetal ActoMyosin Flow (AMF) directed into the contact site. Indeed, RHOA pathway inhibitors significantly reduced Dectin-1 associated DC-SIGN recruitment to the contact site. We used agent based modeling to predict DC-SIGN transport kinetics with (“Directed+Brownian”) and without (“Brownian”) the hypothesized actomyosin flow-mediated transport. The Directed+Brownian transport model predicted a DC-SIGN contact site recruitment (108.72%), similar to that we observed experimentally under receptor co-expression. Brownian diffusive transport alone predicted contact site DC-SIGN recruitment of only 54.02%. However, this value was similar to experimentally observed recruitment in cells without Dectin-1 or treated with RHOA inhibitor, suggesting that it accurately predicted DC-SIGN recruitment when a contact site AMF would not be generated. TIRF microscopy of nascent cell contacts on glucan-coated glass revealed Dectin-1 dependent DC-SIGN and F-actin (LifeAct) recruitment kinetics to early-stage contact site membranes. DC-SIGN entry followed F-actin with a temporal lag of 8.35 ± 4.57 seconds, but this correlation was disrupted by treatment with RHOA inhibitor. Thus, computational and experimental evidence provides support for the existence of a Dectin-1/RHOA-dependent AMF that produces a force to drive DC-SIGN recruitment to pathogen contact sites, resulting in improved pathogen capture and retention by immunocytes. These data suggest that the rapid collaborative response of Dectin-1 and DC-SIGN in early contact sties might be important for the efficient acquisition of yeast under flow conditions, such as those that prevail in circulation or mucocutaneous sites of infection.

## 1. Introduction

C-type lectin receptors play an important role in recognition of the two major fungal cell wall polysaccharide ligands exposed at the host-pathogen interface. DC-SIGN recognizes abundantly exposed N-mannan in the outer cell wall whereas Dectin-1 recognizes nanoscale exposures of β-(1,3)-glucan. Recognition of β-glucan by Dectin-1 contributes to phagocytosis, oxidative burst, regulation of transcription, production of inflammatory cytokines and chemokines, and initiation of adaptive immunity[1]. DC-SIGN is known to mediate intercellular adhesion, as well as antigen uptake and signaling in dendritic cells (DCs)[2]. We are examining the relationship between Dectin-1 and DC-SIGN to understand, in a simplified model, how an effective host-pathogen contact is built. We focused on the earliest events in fungal contact site biogenesis.

Initial pathogen capture and formation of a stable contact site are the earliest events that must occur for signaling through antifungal receptors to initiate. Our previous work with zymosan particles demonstrated that human monocyte derived dendritic cells (DC) form durable contacts between the DC plasma membrane and extracellular fungal particles, which may be important for the antigen gathering functions of these cells [3]. Rapid formation of adhesive contact site structures is especially important for *C. albicans* capture under conditions involving fluid shear stress, for example by reticuloendothelial macrophages capturing yeast in the bloodstream. Fungal recognition under fluid shear also pertains to phagocytes interacting with *Candida* in the oropharyngeal cavity, a major site of mucocutaneous candidiasis, where the host-pathogen interaction is subject to salivary flow.

Various authors have described the accumulation of pattern recognition receptors, such as Dectin-1 and DC-SIGN, at fungal contact sites[4–6]. Immune cells must mobilize receptors to these contact sites for activation, crosstalk and amplification of signaling that directs downstream immune responses. In fact, these contact sites achieve an ordered segregation of molecular components with a peripheral zone enriched in the large transmembrane phosphatase CD45 and a central zone where DC-SIGN and Dectin-1 concentrates. Such “phagocytic synapses” can also involve the development of barriers to molecular diffusion that support specialized signaling processes occurring therein[7,8]. These findings suggest that PRRs are recruited to fungal contacts in some fashion to support their enrichment at these sites. Active and passive transport processes might conceivably account for observed receptor recruitment, but the molecular mechanisms of innate immunoreceptor recruitment in contact sites with *C. albicans* have not been defined.

Previous studies from our group and others have shown the enrichment of DC-SIGN and CD-206 at fungal contact sites[4–6,9]. These studies are typically conducted at longer time scales of hours, which is relevant to processes such as cytokine response and cytotoxic effector responses. However, there is much less information on the dynamics of pattern recognition receptors at fungal contact sites on the time scale of minutes—a time scale that is relevant to the earliest signaling events necessary for innate immune fungal recognition. In the intensely studied immunologic synapse, it is known that immunoreceptors in the T cell/Antigen Presenting Cell (APC) immune synapse are actively transported into the synapse within minutes via their coupling to a centripetal RHOA/Myosin II dependent actomyosin flow (AMF)[10]. Likewise, we previously demonstrated that that Dectin-1 stimulation by glucan activates mechanical contractility signaling via a RHOA/ROCK/Myosin II signaling module within minutes post-stimulation [11]. Thus, the central hypothesis tested in this study is that Dectin-1 activates a transport mechanism, through RHOA/ROCK/Myosin II dependent signaling processes, which facilitates the recruitment of DC-SIGN to the contact site. This would be expected to improve fungal particle retention by providing higher avidity adhesive interactions with the fungal cell wall.

We used a micropipette-micromanipulation approach to provide very high spatiotemporal control over host-pathogen contact site formation. We report that Dectin-1, in collaboration with DC-SIGN, does promote improved capture of *C. albicans* yeast. This occurs through improved recruitment of DC-SIGN to the contact site in a manner that is dependent upon Dectin-1 signaling via RHOA, ROCK and myosin II. These findings provide a high-resolution view of early events in receptor recruitment processes that tailor the earliest stages of the innate immune anti-fungal response.

## 2. Materials and Methods

### 2.1 Cell culture

HEK-293 cells (ATCC, #CRL-1573) were cultured in DMEM containing 10% FBS, 1% penicillin-streptomycin, 2 mM L-glutamine and 1 mM sodium pyruvate at 37°C, in a 5% CO2 environment in an incubator. The identity and mycoplasma-free status of the cell line was independently confirmed by submission to ATCC Human Cell Line authentication (STR) and mycoplasma detection (PCR) services.

### 2.2 Transfection

mApple-Dectin1A-C-10 was a gift from Michael Davidson (Addgene plasmid # 54883; http://n2t.net/addgene:54883; RRID:Addgene_54883). pEGFP-DC-SIGN was a generous gift from Ken Jacobson[12]. pUNO1-hDectin-1a was purchased from Invivogen (#puno1-hdectin1a). mCardinal-Lifeact-7 was a gift from Michael Davidson (Addgene plasmid #54663). Transfection with plasmid was performed following standard protocols for Fu gene 6 (Promega, #E2691). Cells were selected for stable expression using Geneticin (G418 sulfate) (Thermo Fisher Scientific, 10131035; for pEGFP-DC-SIGN) at 400 μg/ml or Blasticidin (Invivogen #ant-bl-05; for pUNO1-hDectin-1a) at 20 μg/ml for 2 weeks.

### 2.3 Micropipette

1.5 mm outer diameter and 1.12 mm inner diameter borosilicate glass capillaries were purchased from World Precision Instruments (WPI# TW150-4). We optimized fabrication procedures to obtain micropipettes with 2 μm diameter tips. We used a microelectrode puller (WPI #micrPUL-1000) for pulling micropipettes. Our final protocol for pulling micropipette of 2 μm was as follows.

**Table 1.** Values of each parameter in WPI microelectrode puller used to pull 2 μm diameter tip micropipettes.

|  | Heat | Force | Distance | Delay |
| --- | --- | --- | --- | --- |
| 1 | 680 | 200 | 2.50 | 55 |
| 2 | 600 | 170 | 1.50 | 60 |
| 3 | 550 | 150 | 0.60 | 35 |
| 4 | 500 | 100 | 0.30 | 20 |

### 2.4 Fungal culture

*C. albicans* clinical isolate TRL035 was obtained as previously described[13]. Isolate was stored as single-use glycerol stock aliquots −80°C. This stock was transferred to 5 ml sterile yeast extract-peptone-dextrose (YPD) medium (Becton Dickinson) at concentration of 1 x 10^5^ cell/ml of YPD and grown for 16 hours at 30°C, with a shaking speed of 300 rpm. The glycerol stock contained 4×10^7^ yeast/ml and was previously calibrated to provide 3 x 10^8^/ml yeast cells at the late log phase under the stated growth conditions.

### 2.5 Silicone chambers

We used silicone isolators (Grace Bio-labs # 665116 and # 665203) on cover glass in configurations as shown below (Fig. 1). Whole chamber was sterilized by passing it through a Bunsen burner blue flame 5 times.

**Figure 1.**
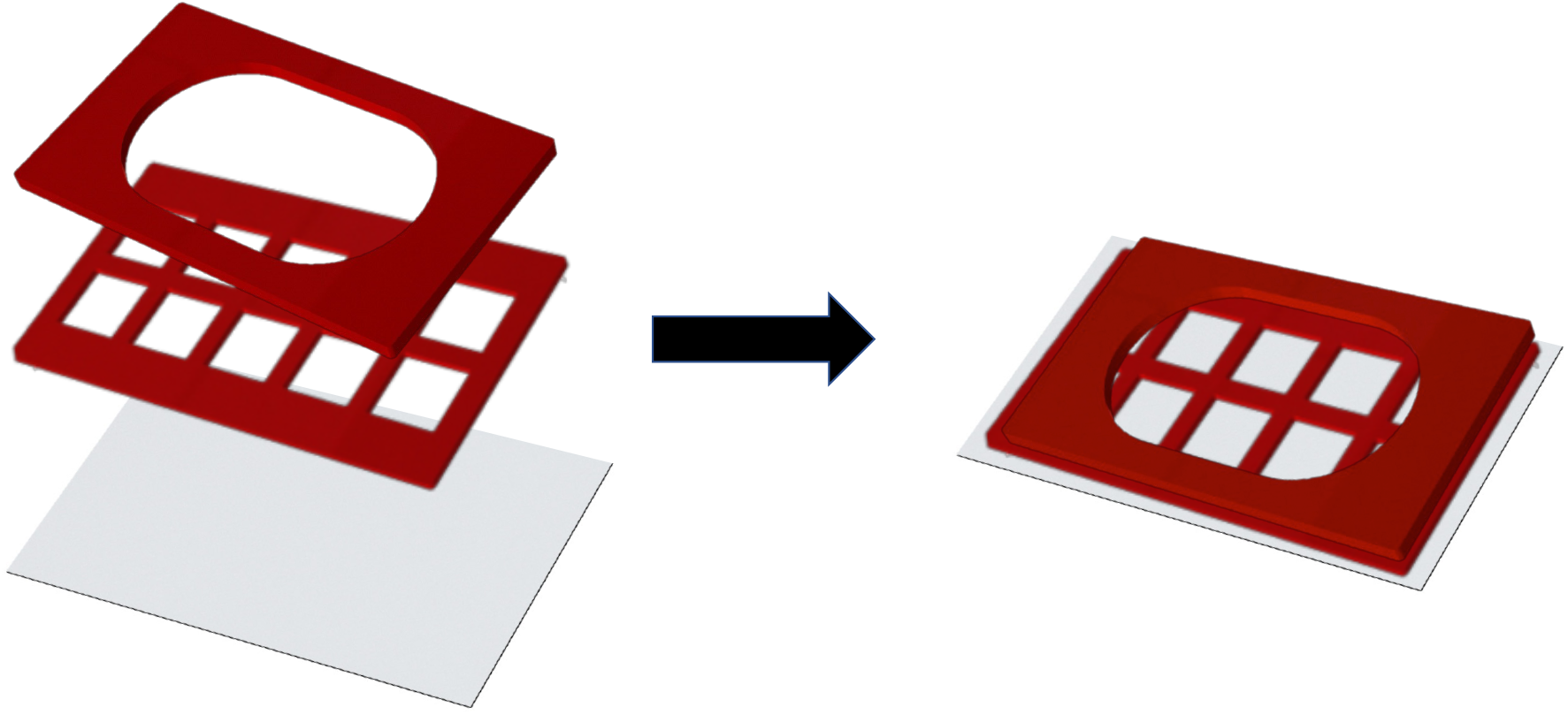
Arrangement of silicone chambers for growing HEK-293 cells for contact site studies.

### 2.6 Contact site studies

HEK-293 cells were transfected with mApple-Dectin1A-C-10 and stable lines were generated, as described above. 2 days before each experiment mApple-Dectin1A-C-10 stable line cells were transiently transfected with pEGFP-DC-SIGN. Next day cells and yeast were seeded into separate compartments of sterilized silicone chambers in a configuration as shown below (Fig. 2).

**Figure 2.**
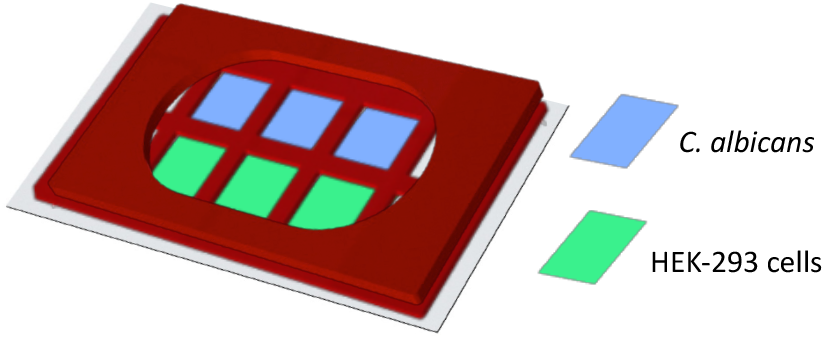
Separation of HEK-293 cells and yeast within chamber.

Cells were stained with CellMask Deep Red Plasma Membrane stain (CMDR) (Invitrogen, #C10046). Original stock CMDR was freshly diluted 1:100 in culture medium to form a working stock solution. 10 μl working stock CMDR was added per 1 ml of culture medium in the culture vessel. Depending on the condition, along with CMDR, various inhibitors were added during this stage. Blebbistatin (Sigma-Aldrich, #203390) at 12.5 μM, or Y-27632 (Sigma-Aldrich, #Y0503) at 5 μM were added for 1 hour. For RHO inhibitor conditions, cells were treated with C3 transferase (Cytoskeleton Inc., #CT04) 1.5 μg/ml for 2 hours. Both CMDR and inhibitor were added to culture dish at the same time. TRL035 *C. albicans* were stained with Fluorescent Brightener 28 (Calcofluor White) (Sigma-Aldrich #F3543). 25 μl of 1 mg/ml Calcofluor White was used to stain 1 ml of yeast in PBS pH 7.4 for 15 min. Then yeast were washed three times with PBS and vortexed for 15 minutes. *C. albicans* were added to smaller chambers as shown in Figure 2. Glucose oxidase (Sigma-Aldrich # G2133) and catalase (Sigma-Aldrich # C100) were added to the chamber at concentration of 0.5 mg/ml and 40 μg/ml, respectively, during data acquisition to reduce photobleaching.

Micropipettes were filled with PBS using MicroFil (WPI #MF28G-5) and syringe. The PBS-filled micropipette was then attached microelectrode holder (WPI # MPH415). The inlet of the microelectrode holder was attached to 1 ml syringe using Luer lock via plastic tubing.

For micromanipulation, we used a Sensapex micromanipulator. We attached microelectrode holder to micromanipulator using an electrode handle (WPI #2505). The silicon chamber was then placed on the FV1000 laser-scanning confocal microscope (Olympus, Center Valley, PA) with controlled temperature, 37°C at 5% CO2. We used a 60x super-corrected, 1.40 NA, Plan-Apochromat oil immersion objective for imaging cells. We replaced one microscope eye piece with a Centering Telescope eyepiece (Olympus # U-CT30-2), which allowed a separate focal plane in each eyepiece for ease in positioning the micropipette.

We identified a suitable single *C. albicans* yeast for capture and then adjusted the microscope’s focal plane to ~20 μm above that yeast. We focused the Centering Telescope eyepiece on the tip of micropipette. The telescope eyepiece was then focused at a plane closer to the yeast, then we lowered the tip of micropipette via micromanipulator into the lower focal plane toward the yeast, while being monitored through the telescopic eyepiece. This process was repeated till we reached the level of fungus. This was done to avoid breaking of micropipette tip while lowering it. We always lowered telescopic eyepiece focus first and then adjusted micropipette level. Once at the same level as *C. albicans,* we applied negative pressure using the syringe to capture a single yeast on the micropipette tip. Then, we manipulated *C. albicans* to the chamber with HEK-293 cells by moving from the yeast chamber, over the isolator barrier, and translating into the cell chamber, keeping the tip submerged at all times. Finally, *C. albicans* was brought near the plasma membrane of a HEK-293 cell expressing the appropriate receptors, as verified by their fluorescent protein tags.

Confocal fluorescence microscopic observation of contacts sites was conducted with the following parameters. Calcofluor White (a marker for all yeast) was excited with a 50 mW, 405 nm diode laser operated at 1% power, and CMDR was excited with a 20 mW, 635 nm diode laser operated at 0.5% power. EGFP–DC-SIGN was excited with a 20 mW, 473 nm diode laser operated at 1% power. mApple-Dectin-1 was excited with a 20 mW 559 nm laser at 1% power. These lines were reflected to the specimen by a 405/473/559/ 635 multi-edge main dichroic element and routed through a confocal pinhole (110 mm diameter) to secondary dichroic followed by bandpass emission filters in front of two independent PMT detectors and two independent high-sensitivity GaAsP PMT detectors (HSD). Specifically, the emission light passed by the main dichroic was directed to PMT1 (Calcofluor White channel) via reflection from the SDM473 dichroic and passage through a BA430-455 nm bandpass filter. For the CMDR channel, light from SDM473 was directed to a 640 nm shortpass dichroic and BA575-675 nm bandpass filter. Light from 640 nm shortpass dichroic was directed to SDM560 filter cube to HSD1 (the EGFP–DC-SIGN channel) via passage through a BA490-540 nm bandpass filter. For mApple-Dectin-1, light was directed via SDM560 filter cube to HSD2 via passage through a BA575-675 nm. 60x lens with 3x zoom was used to capture images. Further a subregion of interest for image scanning was selected such that the region was small enough to be scanned at a rate of 0.400 sec per frame. Overall, pixel size for all images was 7.24 pixels per micron. Imaging was started and then, with the micromanipulator, contact was made between *C. albicans* and HEK-293 cells. The contact site was imaged for 10 minutes total duration after contact initiation.

### 2.7 Polystyrene bead control for contact site studies

For making Dextran coated beads, we used 5 μm streptavidin-coated polystyrene bead (Spherotech #VP-60-5). We used 1,1’-carbonyldiimidazole in DMSO based system as described by Tam et al. to conjugate dextran (Sigma-Aldrich # 31388) with beads. Rest of the procedure for making contact and imaging was exactly same for TRL035 *C. albicans*.

### 2.8 FRAP studies

For FRAP studies, exact same steps as mentioned above for contact site studies were followed. Then TRL035 contact site was allowed to mature for 10 min. after contact. Then a rectangular FRAP window was selected so that it included the whole of the contact site. Imaging was started and 5 frames were collected pre-bleach. Then, we photobleached the contact area with 473 nm and 559 laser, 100% power for 500 milliseconds. Imaging was continued for 10 min. to quantify recovery. FRAP analysis was done using easyFRAP[14].

### 2.9 Analysis of contacts site data

For quantifying the contact site, we used the Fiji distribution of ImageJ. We demarcated overlapping pixels between dilated calcofluor channel and CMDR channel. These overlapping pixels denote the contact site. All further calculation of MFI (Mean Fluorescence Intensity) for DC-SIGN, Dectin-1 channel and their normalizations were done from these contact site pixels only. The detailed steps followed were as follows. Calcofluor white (405 channel) was thresholded to make a fungal mask and converted to binary. The binary fungal mask was then dilated 2 times. The fungal mask was divided by 255 to make all pixel values 1. This is essential for calculating overlapping areas in next steps. The CMDR (635 channel) was thresholded and converted to binary to create a CMDR mask. The CMDR mask was divided by 255. The fungal mask was multiplied by CMDR mask to mark overlapping pixels as those demarcating the contact site mask, within which each pixel had a numerical value of 1. The remaining non-mask pixels had a numerical value of 0. Overlapping pixels were multiplied with Dectin-1, DC-SIGN and CMDR raw pixel intensities, creating masked Dectin-1, DC-SIGN and CMDR datasets. RawDensity, which is the sum of intensities of all pixels in a dataset, was calculated for each the contact site masked DC-SIGN, Dectin-1 and CMDR datasets. The same calculation was performed for the contact site mask image, which provides the area (pixels) of the contact site mask. Mean Fluorescence Intensity (MFI) per pixel for DC-SIGN, and Dectin-1, and CMDR were calculated by dividing RawDensity for each of these datasets by the contact site area in pixels. To normalize for variable amount of membrane in a contact site, we divided MFI of DC-SIGN and Dectin-1 by the corresponding contact site CMDR MFI. Finally, to control for possible differential expression/staining of individual HEK-293 cells, we expressed the above normalized receptor MFI signals as a percentage of their value at time 0, on a per cell basis.

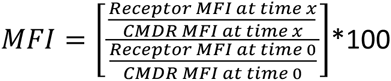

### 2.10 Yeast capture assay

For the yeast capture assay, we used HEK-293 cells stably transfected mApple-Dectin1A-C-10 cells were used or transiently transfected with pEGFP-DC-SIGN. For DC-SIGN only condition parental cells were transiently transfected with pEGFP-DC-SIGN. Overall, the following 4 conditions were used for experiments, 1) EGFP-DC-SIGN + mApple-Dectin-1, 2) EGFP-DC-SIGN, 3) mApple-Dectin-1, 4) Untransfected conditions. Cells were seeded at 2.5 x 10^4^/dish, overnight in 35 mm Mattek dishes. Then TRL035 *C. albicans* stained with calcofluor white were added to dishes at 10 x 10^4^/dish. Then dishes were kept on rocker shaker (BR200 2d rocker, Southwest Science) for 30 min. Cells were then washed 3 times with PBS and fixed with 4% PFA, and the number of fungi attached to each cell of interest were counted under microscope with imaging condition similar to described in contact site studies. Binding studies with cells expressing both DC-SIGN and Dectin-1 enumerated yeast binding only to cells confirmed to co-express both receptors.

### 2.11 TIRF microscopy

HEK-293 cells were stably transfected with pUNO1-hDectin-1a and mCardinal-LifeAct-7. 1 days before experiment these cells were transiently transfected with EGFP-DC-SIGN. Cells were dissociated from dish surfaces using brief exposure to 0.25% trypsin-EDTA followed by addition of protein rich medium to rapidly quench trypsin activity and washing into fresh culture medium. Then these cells were put on 35 mm dish coated with β-glucan. Cells were allowed to settle down on these surfaces in incubator for 30 min. Then cell membrane was observed under Olympus IX83 TIRF/Single Molecule Localization Microscope. 488 nm and 561 nm lasers were used to excite EGFP-DC-SIGN and mCardinal-LifeAct respectively. Cell membrane was observed for 5 min. To address the potential concern that trypsinization would damage transmembrane receptors sufficiently to render cells non-responsive to fungal ligands, we loaded cells with Fluo-4 calcium dye, trypsinized and settled them on glucan coated glass as above, and observed calcium flux upon Dectin-1 contact with glucan coated glass. We observed calcium flux on glucan coated glass but not on glucan free glass surfaces, demonstrating that the cells remained functional following trypsinization (data not shown)

### 2.12 Coating dishes with β-glucan

The central glass region of Mattek glass bottom 35 mm dishes was coated with β-glucan to permit TIRF microscopic observations of early contact site membrane dynamics using a coupling chemistry previously reported by Tam, et al.[6]. The following procedure was used to produce these surfaces. 200 μl of 0.01 w/v Poly-l-lysine aqueous solution (Sigma, P4707) was put on the central portion of 35 mm dishes and adsorption was permitted to occur for 30 min. Excess solution was removed and dishes were washed with DMSO for 3 times. Poly-l-lysine amines were activated by immersion in 0.5M 1,1′-Carbonyldiimidazole (CDI; Sigma, 115533) in DMSO for 1 hour. Dishes were washed with DMSO 3 times. 10 mM Medium Molecular Weight (145 kDa) β-(1,3)-glucan from ImmunoResearch Inc. (Eagan, MN) in DMSO was added to the dishes. The reaction was allowed to incubate overnight, excess solution was removed, and dishes were washed with water 3 times.

### 2.13 Phagocytosis assay

TRL035 *C. albicans* were first stained with Fluorescent Brightener 28 (calcofluor white) (Sigma-Aldrich #F3543). 25 μl of 1 mg/ml calcofluor white was used to stain 1 ml of 16 hour fungal culture in PBS (Gibco) pH 7.4 for 15 minutes. Then cells were washed 3 times with PBS and stained with 75 μM CypHer5E NHS ester (GE Healthcare, PA #15401) for 1 hour at 25°C[13]. *C. albicans* were then added to culture dishes with HEK-293 cells transiently transfected with pEGFP-DC-SIGN for 1 hour and observed under a microscope for increased fluorescence in CypHer5E channel as an indicator of fungi which had been phagocytosed.

### 2.14 Agent Based Modeling

<u>An agent-based model of DC-SIGN transport into contact sites was created in the Netlogo modeling environment. Source code and a full description of the parameterization of this model are provided in Supplemental Methods</u>.

## 3. Results

### 3.1 Dynamics of Dectin-1 and DC-SIGN recruitment to a contact site for capture of C. albicans

We examined the possible role of Dectin-1 and/or DC-SIGN in the capture of fungal particles using a *C. albicans* yeast capture assay. This assay was designed to test the ability of cells expressing these receptors, alone or in combination, to bind and stably capture yeast after a relatively brief exposure to particles under conditions of fluid shear. To unambiguously measure fungal capture attributable to these receptors, we used a HEK293 cell system with no endogenous expression of relevant PRRs, which was then transfected to express one or both receptors. We found that, within 30 min., cells captured an average of 5.1, 1.62, 2.23 and 0.23 *C. albicans* fungal particles per cell under DC-SIGN + Dectin-1, Dectn-1 only, DC-SIGN only and untransfected/parental HEK-293 conditions, respectively (Figure 3a). We found significantly increased capture of fungal particles in the co-expressed condition compared to the condition with either individual receptor expressed alone. This effect was slightly more than additive. Given the normal distribution of yeast capture measured in co-expressing cells, we found that there is only 3.2% probability to find a fungal capture equivalent to the sum of means of yeast capture measured for cells expressing DC-SIGN and Dectin-1 alone. We hypothesized that DC-SIGN and Dectin-1 collaborate to improve capture of *C. albicans* yeast in rapidly developing contact sites with this pathogen.

**Figure 3.**
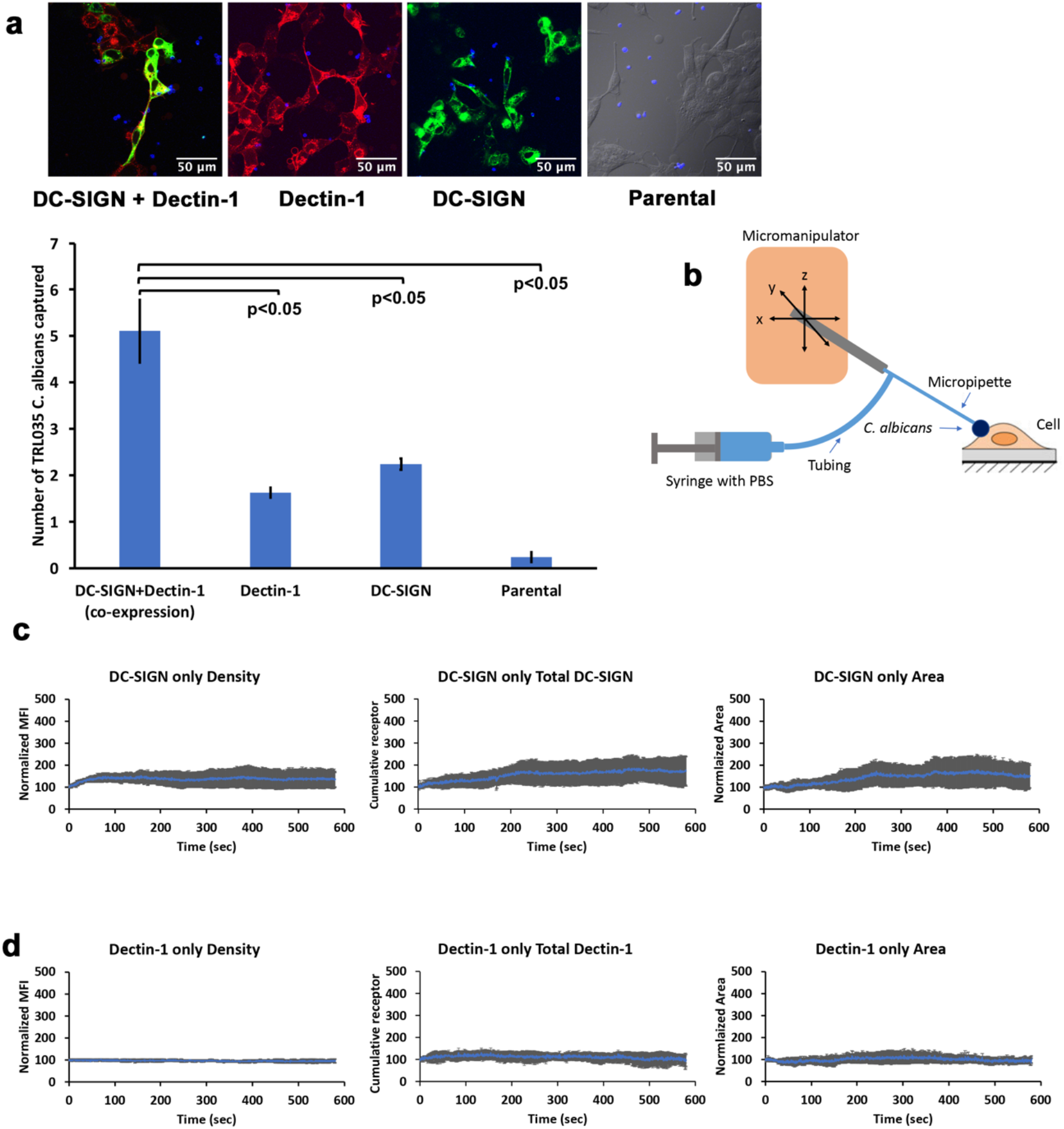
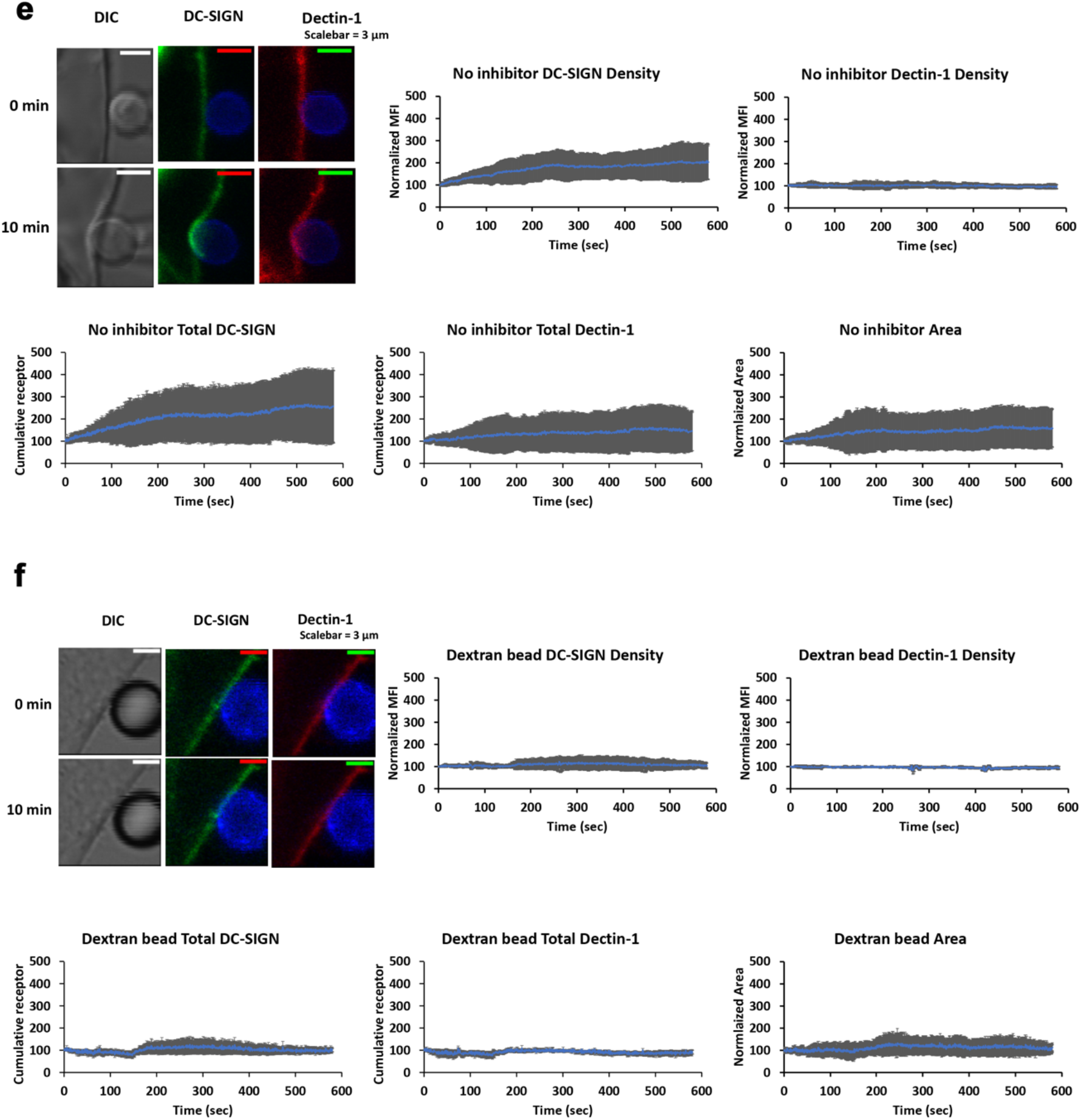
Dynamics of DC-SIGN and Dectin-1 recruitment to a contact site for capture of *C. albicans*. **(a)**TRL035 *C. albicans* (blue) capture assay by HEK-293 cells transfected with EGFP-DC-SIGN (green) and/ mApple-Dectin-1 (red) (n=61), Dectin-1 only (n=63), DC-SIGN only (n=43) and parental (n=49). Average number of fungal particles captured by a cell under each condition is denoted in the graph (the reported value of “n” denotes the number of total cells pooled from ≥3 independent experimental replicates). **(b)** Schematic diagram of micropipette-micromanipulator system. **(c & d)** Dynamics of DC-SIGN (panel c, n=6) or Dectin-1 (panel d, n=6) density enrichment at the contact site under individual receptor expression conditions. **(e)** Images showing contact of TRL035 *C. albicans* (blue) with HEK-293 cells transfected with EGFP-DC-SIGN (green) and mApple-Dectin-1 (red) at time 0 and time 10 min. Graphs show dynamics of normalized DC-SIGN and Dectin-1 density enrichment, total cumulative receptor intensity and contact site area under co-expression condition of receptors at TRL035 contact site (n=10). **(f)** Dynamics of EGFP-DC-SIGN (green) and mApple-Dectin-1 (red) density enrichment under co-expression of receptors at contact site with dextran coated polystyrene bead (blue) (n=3). All graphs represent the mean ± S.D. for the indicated value at each time point).

To look for receptor recruitment dynamics at the contact site, we used micropipette-micromanipulator based system where a single fungal particle was adhered to the tip of the micropipette and advanced into contact with the cell to form a single host-pathogen contact site at a well-defined location and time (Figure 3b). Fluorescent protein tagged receptors’ recruitment to the contact site was followed by live cell confocal fluorescence microscopic observation of the contact site from initiation to 10 minutes post-contact and reported as a relative increase over receptor density at the time of contact site initiation. We additionally reported changes in cumulative receptor intensity (sum of all pixel intensities in contact) and contact site area. First, we measured recruitment of EGFP-DC-SIGN or mApple-Dectin-1A individually expressed in HEK293 cells to contact sites formed as described above. In the DC-SIGN only condition, we found a 35.04% increase in normalized density, 70.76% increase in cumulative receptor intensity and 49.83% increase in the contact site area (Figure 3c). For Dectin-1 only expression, we found no significant increase in normalized Dectin-1 density, cumulative receptor fluorescence and contact site area (Figure 3d). Then, we examined contact sites in HEK-293 cells co-transfected with both EGFP-DC-SIGN and mApple-Dectin-1A (Figure 3e). In dual expressing cells, we found a significant increase in normalized DC-SIGN density of 103.15% compared to pre-contact DC-SIGN density. We did not find a significant increase in Dectin-1 density within 10 minutes post contact in dual expressing cells, similarly to the observation in Dectin-1 only expressing cells. For total amount of receptor in contact sites, we found 154.15% and 45.33% increase in cumulative receptor intensity for DC-SIGN and Dectin-1 respectively. We also found 57.97% increase in contact area in dual expressing cells. With control dextran coated polystyrene beads, we did not find any increase in DC-SIGN or Dectin-1 recruitment (Figure 3f). In summary, we found that DC-SIGN exhibited significantly increased recruitment and accumulation at contact site in the presence of Dectin-1 within the 1^st^ 10 min of *C. albicans* contact. Whereas, Dectin-1 did not show recruitment within the 1^st^ 10 min of *C. albicans* contact.

### 3.2 Actomyosin based active recruitment of DC-SIGN at C. albicans contact site

We previously reported a novel signaling connection between Dectin-1 activation and generation of cellular mechanical forces via stimulation of actomyosin contractility [11]. Actomyosin contractility has previously been found to generate actomyosin flows (AMFs) that support active transport of plasma membrane proteins (i.e., TCR and BCR transport at immunological synapses, [10]). Therefore, we next looked for possible involvement of the Dectin-1 activated actomyosin contractility signaling mechanism in early active recruitment of DC-SIGN to fungal contact sites. We hypothesized that Dectin-1 mediated actomyosin contraction leads to an AMF at host-pathogen contact sites, and that DC-SIGN coupling to this AMF is important for recruitment of DC-SIGN to *C. albicans* contact sites. This is feasible because Dectin-1 signaling gives rise RHOA-ROCK-Myosin II activation, leading to actomyosin contractility[11]. So, inhibition of this signaling mechanism would be expected to abrogate the development of a hypothesized AMF and any active receptor recruitment connected to it. We pretreated HEK-293 cells transfected with DC-SIGN and Dectin-1 with various inhibitors of the actomyosin contractility signaling axis. We found that cells treated with inhibitors of RHOA, ROCK and Myosin-II exhibited normalized DC-SIGN density of 41.47%, 62.96% and 44.04% respectively at 10 minutes post-contact formation (Figure 4a). Thus, there was significantly less recruitment of DC-SIGN in the presence of each of the 3 inhibitors relative to the untreated condition. For total cumulative DC-SIGN intensity within the contact sites, we found an increase of 71.13%, 78.79% and 43.03% with RHOA, ROCK and Myosin-II inhibitor (Figure 4b). Thus, showing less total accumulation of DC-SIGN in inhibitor treated contacts as well. Concomitantly, we observed contact site areas increased by 121.77%, 105.43% and 18.36% with RHOA, ROCK and Myosin-II inhibitor, respectively, at 10 minutes post-contact formation (Figure 4c). Thus, there was an increase in contact area with RHOA and ROCK inhibitor, but a decrease in contact area with Myosin-II inhibitor compared to no inhibitor condition.

**Figure 4.**
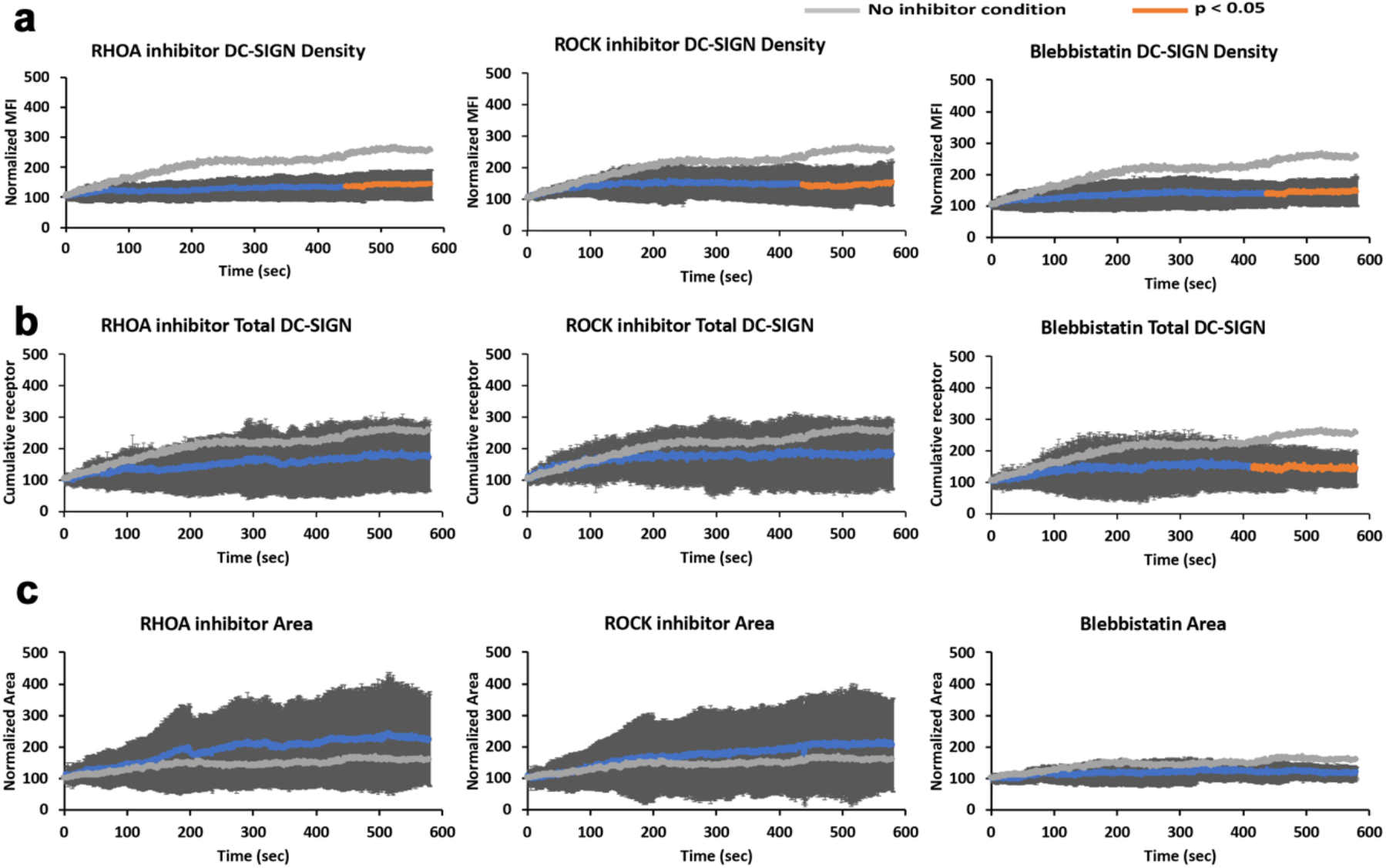
Role of RHOA-ROCK-Myosin II based system in recruitment of DC-SIGN under co-expression condition of Dectin-1 and DC-SIGN at TRL035 *C. albicans* contact site. **(a)** Normalized receptor density enrichment in fungal pathogen contact sites for DC-SIGN at the end of 10 min. under RHOA (n=11), ROCK (n=9) and Myosin-II (n=9) inhibitory conditions, with comparison to the same in untreated cells. **(b)** Total cumulative DC-SIGN intensity enrichment under the same conditions as panel a. **(c**) Contact site area changes under the same conditions as panel a. All graphs represent the mean ± S.D. for the indicated value at each time point.

### 3.3 Diffusivity of DC-SIGN and Dectin-1 within contact site

Having examined the mechanisms of Dectin-1 dependent DC-SIGN mobilization to contact sites, we finally examined the mobility of these receptors once they arrived at the contact site. We hypothesized that receptor engagement with fungal cell wall ligands after transport into the contact site would result in reductions in apparent receptor lateral mobility, especially for DC-SIGN due to its multivalent binding to cell wall mannoproteins. We used FRAP (Fluorescence Recovery After Photobleaching) on contact sites after 10 minutes of contact site maturation (Figure 5a). Within contact sites, we found that DC-SIGN showed 28% mobile fraction with recovery t_1/2_ of 66.19 sec and Dectin-1 showed much larger 79% mobile fraction with recovery t_1/2_ of 42.13 sec (Figure 5b). On statistical comparison of this contact site FRAP data we found that, relative to Dectin-1 values, DC-SIGN had lower lateral mobility (p=0.01) and exhibited a smaller mobile fraction (p=0.0003). For non-contact membrane, we found that DC-SIGN showed 78.82% mobile fraction with recovery t_1/2_ of 35.13 sec and Dectin-1 showed 79.66% mobile fraction with recovery t_1/2_ of 14.08 sec (Figure 5c). On statistical comparison of DC-SIGN within contact sites with non-contact DC-SIGN, we found a significant increase in recovery t_1/2_ (p=0.005) and significant decrease in mobile fraction (p=0.0002). For Dectin-1, we found significant decrease in recovery t_1/2_ compared to non-contact membrane (p=0.001) and no significant difference for mobile fraction (p=0.39). These findings indicate that DC-SIGN is far less mobile than Dectin-1 within contact sites. This conclusion is consistent with the expectation that highly multivalent DC-SIGN/N-mannan interactions are likely to result in largely irreversible adhesive interactions of DC-SIGN nanodomains with the outer cell wall surface of *C. albicans*—the likely physical basis of this receptor’s importance for rapid and effective capture of yeast. In contrast, Dectin-1 is primarily monovalent [15] as it enters the contact site, and sites of glucan exposure are quite small and sparsely distributed [16]. Thus Dectin-1 remains fairly mobile within the contact site, which may increase the efficiency of its search for rare sites of glucan exposure even though t_1/2_ is decreased in the contact as expected from Dectin-1 interactions with available glucans at contact sites.

**Figure 5.**
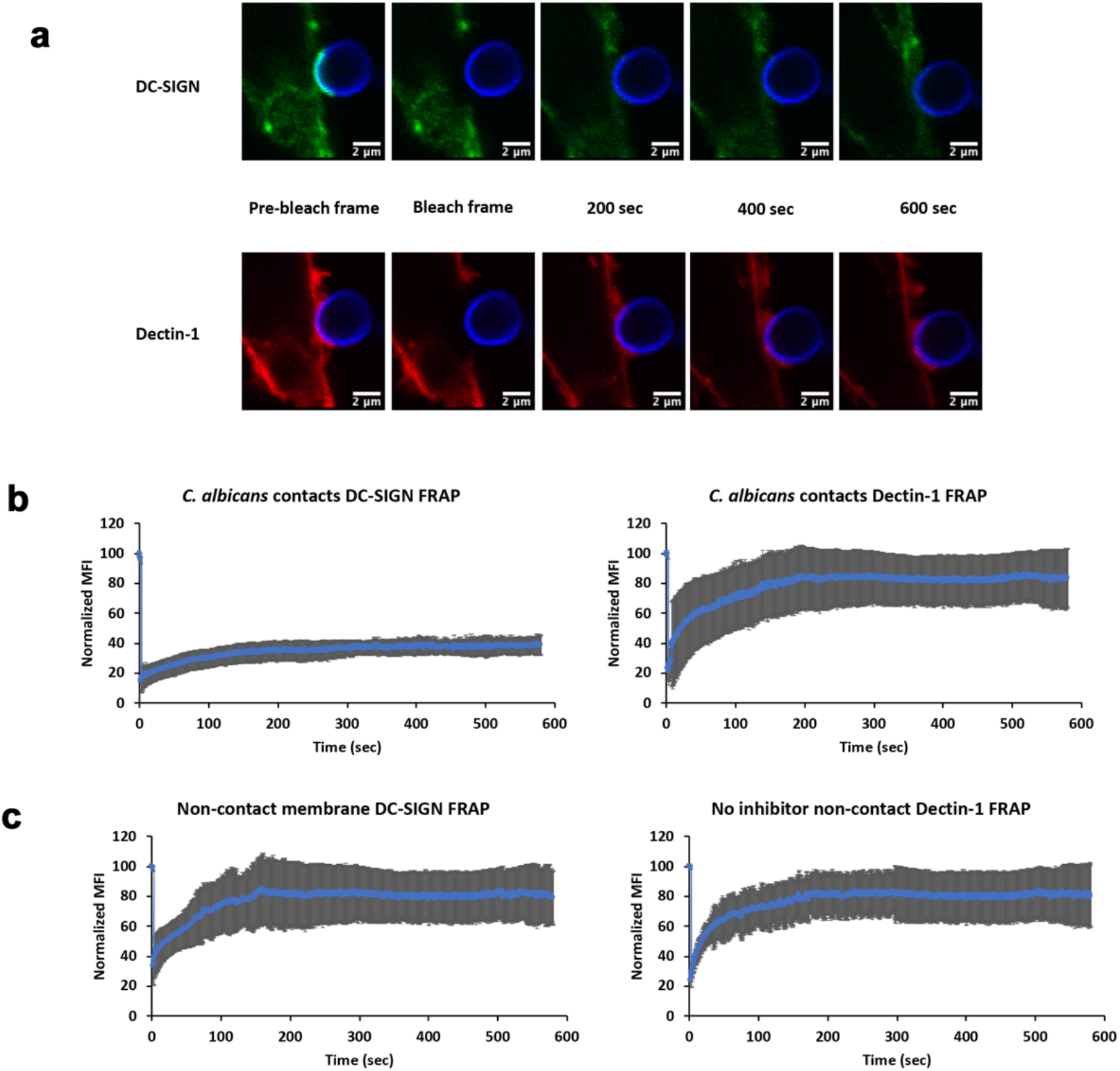
Lateral mobility characteristics of DC-SIGN and Dectin-1 at *C. albicans* inside and outside of contact site zones. **(a)** Example pre/post-FRAP images of TRL035 *C. albicans* (blue) contacts with HEK-293 cells transfected with EGFP-DC-SIGN (green) and mApple-Dectin-1 (red). **(b)** Mean FRAP recovery curves of DC-SIGN (n=7) and Dectin-1 (n=7) at contact sites and **(c)** non-contact membrane DC-SIGN (n=6) and Dectin-1 (n=6) with fluorescence recovery observed for 10 min. All contact sites were allowed to mature for 10 min. after *C. albicans* contact before initiation of FRAP experiments. All graphs represent the mean ± S.D. for the indicated value at each time point.

To rule out effects of RHOA inhibitor itself on receptor lateral mobility, we did FRAP studies on cells exposed to RHOA inhibitor (Figure S2). These cells did not have fungal contact sites to facilitate comparison of effects on basal receptor mobility of the receptors. On comparing DC-SIGN lateral mobility in RHOA inhibited cells to untreated controls, we found no significant difference in DC-SIGN recovery t_1/2_ (p=0.66) or mobile fraction (p=0.71). The same was true for Dectin-1, with no significant difference in t_1/2_ (p=0.53) or mobile fraction (p= 0.22). We conclude that RHOA pathway inhibition does not impact the lateral mobility of DC-SIGN or Dectin-1, making such a direct effect an unlikely contributor to the recruitment of these receptors to the contact site.

### 3.4 Evidence for active transport of DC-SIGN into the C. albicans contact site

In resting cell membranes, the DC-SIGN tetramer self assembles into nanodomains, which have been described to undergo Brownian diffusion as well as apparent directed transport in some contexts[17–19]. For instance, DC-SIGN nanodomain tracking studies have reported that this receptor can undergo rapid, linear transport in the plane of the plasma membrane, suggesting that some directed mobility of DC-SIGN is possible[12,18,20,21]. As previously mentioned, directed transport at immunological contact sites (e.g., T cell or B cell synapses) often involves cytoskeleton/motor protein dependent centripetal motion of receptors toward the center of the contact site[10]. So, we proposed that similar mechanism might exist at the fungal contact site. Therefore, we next used an agent based model to test the hypothesis that a combination of Brownian diffusion and periods of directed transport into the contact site could explain observed DC-SIGN recruitment dynamics (“Directed + Brownian transport” model).

We created an agent-based computational model in the Netlogo modeling environment, which was capable of predicting DC-SIGN accumulation rates in contact sites wherein the DC-SIGN experienced a variable coupling to a centripetally directed active transport mechanism (i.e., AMF), interspersed with regimes of Brownian diffusion when uncoupled to the directed transport mechanism. This model was parameterized with literature reported measurements and systematic parameter studies, as described in Supplemental Methods. The key parameter in this model determining the degree of active transport of DC-SIGN was the AMF coupling coefficient, which we systematically varied to define a range of predicted DC-SIGN contact site recruitment. The Directed+Brownian transport model was able to predict a DC-SIGN contact site recruitment rate of 108.72% increase over pre-contact density (Figure 6), similar to that observed experimentally in cells expressing DC-SIGN and Dectin-1 with 103.15% increase over pre-contact density (Figure 3e). To achieve this density in the model, a coupling coefficient of 40% was required, indicating the strength of DC-SIGN’s attachment to the hypothesized directed transport mechanism. While there is no direct measurement of this value for DC-SIGN available at this time, this coupling coefficient is within the physiological range observed in other systems involving receptor coupling to AMF[10,22]. This result suggests that partial coupling of DC-SIGN to a centripetally directed active transport mechanism is a viable model for explaining experimentally observable DC-SIGN recruitment to *C. albicans* contact sites.

**Figure 6.**
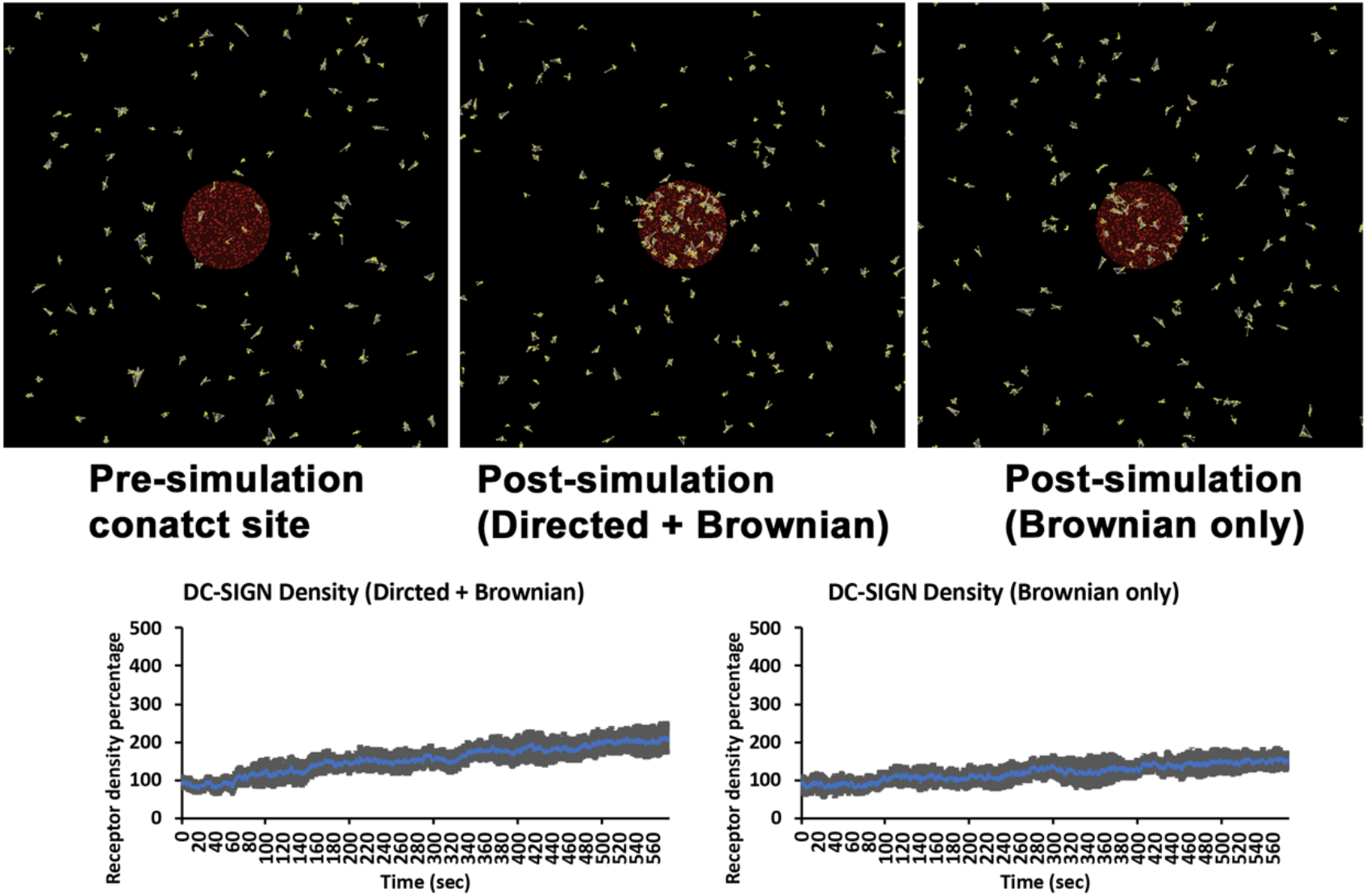
Agent-based modelling showing role of directed motion in DC-SIGN transport at *C. albicans* contact sites. Top panels are views of DC-SIGN (yellow) distribution relative to the contact site (red) before the start (top left) and after 10 minutes of simulation time for models simulating Brownian diffusion only (top right) or directed transport superimposed upon Brownian diffusion (Directed+Brownian) (top middle). Graphs indicate the percentage increase in DC-SIGN density over pre-contact density within 10 minutes of simulation time for Brownian diffusion only model (bottom right) or Directed+Brownian model (bottom left).

To determine the level of DC-SIGN recruitment that could be achieved in the absence of directed motion, we ran simulations at a directed transport coupling coefficient of zero, leaving only lateral mobility via Brownian diffusion (i.e., “Brownian” model). All other parameters of the model were kept exactly the same as in the Directed + Brownian model. We found that the Brownian model predicted contact site DC-SIGN density increase of 54.02% (Figure 6). This is similar to what we observed in inhibitor conditions which are supposed to inhibit directed transport of DC-SIGN at contact site (Figure 4).

If an AMF forms subjacent to the plasma membrane at nascent fungal contacts, we expected an enrichment of F-actin and DC-SIGN in these regions during the early stages of contact site formation. TIRF microscopy is a useful approach with very high axial and lateral resolution for visualizing protein recruitment dynamics at or very near the plasma membrane. However, TIRF microscopy requires the membrane region of interest to be immediately adjacent to the cover glass—a condition not met in our previous contact site experimental model. We therefore devised a contact site model amenable to TIRF microscopy wherein HEK-293 cells transfected with EGFP-DC-SIGN, pUNO1-hDectin-1a and mCardinal-LifeAct-7 were dropped onto β-glucan-coated cover glass surfaces. As cells began to interact with this surface, they extended dynamic protrusive structures to explore the glucan-coated glass. We found that DC-SIGN and F-actin (LifeAct) became enriched within cell processes in contact with the glucan-coated surface, and that the kinetics of this enrichment showed a temporal correlation of DC-SIGN and LifeAct fluorescence, but with a lag of DC-SIGN relative to F-actin recruitment. The observed latency of maximum DC-SIGN intensity relative to LifeAct was 8.35 ± 4.57 seconds. This correlation was almost completely abolished by RHOA inhibitor, with DC-SIGN latency of 1.69 ± 4.97 seconds in this condition (Figure S3), which was a statistically significant decrease (p=0.024) relative to the non-inhibited condition. Thus giving direct imaging based preliminary evidence for Dectin-1 mediated RHOA dependent actomyosin based transport of DC-SIGN at *C. albicans* contact site.

## 4. Discussion

At *C. albicans* contacts with immune cells, which have been described as the “phagocytic synapse”, Goodridge et al. showed the accumulation of Dectin-1 within contact sites between myeloid cell types and model fungal particles and fungal pathogen cells wherein regulatory tyrosine phosphatases CD45 and CD148 were excluded from Dectin-1 rich zones of the contact [7]. The evident recruitment of immunoreceptors at these cellular synaptic structures and alterations to their patterns of lateral mobility in the membrane suggest that it is important to understand the molecular mechanisms responsible for their construction. In the case of fungal host-pathogen contacts, many studies have examined receptor distribution at tens of minutes to hours due to the difficulty of achieving precise control over contact site formation that is necessary for examining the earliest stages of host-pathogen interaction. We have overcome that problem in this study by the use of micropipette-micromanipulator based application of fungal particles to cells with high spatiotemporal precision.

We looked at the dynamics of DC-SIGN and Dectin-1 recruitment at the ≤10-minute time scale. We propose that events occurring at these earliest stages of contact site formation are important to promote pathogen capture and to stabilize the phagocytic synapse. DC-SIGN showed significant contact site recruitment within the first 10 min., whereas Dectin-1 did not show recruitment within this period, relative to its initial density. DC-SIGN is optimized for high avidity interactions with fungal pathogens due to its tetramerization via its stalk domain and its organization into multi-tetramer nanodomains[17,18,23]. Improvements in the efficiency of contact site recruitment of DC-SIGN are likely to be important for determining the ability of the phagocytic synapse to retain a fungal pathogen, especially under circumstances where fluid shear forces could destabilize the contact site before internalization of the particle can take place.

In the absence of Dectin-1, there was a significant decrease in the recruitment of DC-SIGN. In recently published work, we showed that Dectin-1 stimulation by β-glucan gives rise to RHOA-mediated actomyosin activation for contractile mechanical force generation. Previous research on the immunological synapse has demonstrated the importance of RHO-GTPase mediated actin cytoskeleton organization in adhesion and early immunological synapse formation[24]. Further, the work of Tsourkas et al. with B cell synapses showed with stochastic simulations that the formation of the synapse occurs only if BCR mobility is enhanced by directed motor-driven transport [25]. Also, Manzo et al. showed the role of lateral mobility of DC-SIGN nanoclusters in enhancing pathogen binding using Monte Carlo simulations [18]. Because Dectin-1 activation appeared to enhance DC-SIGN recruitment, we tested the role of RHOA-mediated actomyosin activation downstream of Dectin-1, leading to active recruitment of DC-SIGN to *C. albicans* contact sites. Using agent-based modeling, we found that a directed transport process for DC-SIGN recruitment was necessary in order for computational predictions of DC-SIGN recruitment kinetics to match experimentally observed rates of DC-SIGN recruitment to host-pathogen contact sites. To further support this finding experimentally, we found that inhibitors of RHOA, ROCK and Myosin II decreased DC-SIGN recruitment to contact sites (Figure 4). The density of contact site DC-SIGN achieved in the presence these inhibitors was similar to the DC-SIGN density predicted by computational modeling of contact site biogenesis under the assumption that DC-SIGN was only transported by passive diffusion followed by trapping in the contact via high avidity interactions with cell wall N-mannans (Figure 6). This result strongly suggests that RHOA, ROCK and Myosin II are essential components of a Dectin-1 dependent active transport mechanism that enhances DC-SIGN recruitment to the sites of host-pathogen interaction within minutes of pathogen contact.

RHOA and ROCK inhibited contacts exhibited a larger total area than without inhibitor. We expect that the spreading of a cell over a pathogen surface to form a contact site is a net result of protrusive (e.g., RAC1-mediated branched actin in lamellar edges and membrane ruffles) and contractile processes (i.e., RHOA-mediated actomyosin contraction around the contact), both of which are unfolding over the time scale of our studies. The fact that RHOA pathway inhibitors allow greater cell spreading to form larger area contacts is compelling evidence that contractile mechanical forces are being generated during the first ten minutes of host-pathogen interaction with *C. albicans*.

We did not find significant enhancement of Dectin-1 density in contact sites relative to non-contact membrane within the first 10 min. of *C. albicans* contact. We think this is because sites of nanoscopic exposures of β-glucan on the *C. albicans* wall surface are limited in spatial extent and sparsely distributed. As described by Ostrowski et al. for phagocytic synapse, upon initial clustering of cognate receptors the ligands on the pathogen surface trigger the signaling pathways that could initiate signaling and phagocytosis[8]. However, receptor density enrichment and continuous receptor-ligand interaction is required to complete internalization of phagocytic particle, otherwise phagocytosis will stall[26]. Our results are consistent with a model of *C. albicans* recognition wherein initial Brownian diffusion of Dectin-1 leads to rapid Dectin-1 engagement and RHOA-mediated actomyosin flow formation (Figure 6). Subsequently, DC-SIGN gets coupled to this actomyosin flow, which facilitates its efficient mobilization to the contact site. Soon after being drawn into the contact site, DC-SIGN nanodomains engage in high avidity interactions with the fungal cell wall surface, which is important for strong retention of the fungal particle. The high fraction of pathogen-interacting DC-SIGN in the contact site is evident from our FRAP results where the majority of DC-SIGN is immobile over the time scale of minutes (Figure 5). It is this prolonged pathogen retention that allows Dectin-1, the pro-phagocytic receptor, to engage in the more time-consuming process of searching for and integrating signaling from sparse sites of nanoscale glucan exposure. This is evident from a large mobile fraction of Dectin-1 in contacts sites at 10 min. At larger time scales, Dectin-1 may show significant enrichment within phagocytic contacts to amplify its signaling ultimately giving rise to phagocytosis. This is supported by Tam et al. showing accumulation of Dectin-1 within phagocytic synapse with glucan coated particles at 30 min. time scale[6]. Also, Strijibis et al. showed importance BTK, VAV1 and F-actin accumulation within *C. albicans* contact site for efficient phagocytosis of fungus at an hour scale[5]. Our results on the earliest stages of innate immune fungal recognition provide some increased insight into host-pathogen contact site evolution, which is evidently a complex, orchestrated process that involves many receptors being are recruited and activated across different time scales.

Yi et al. showed direct evidence for actin retrograde flow and actomyosin II arc contractions playing role in driving TCR cluster at T-cell immunologic synapse[10]. We hypothesize that DC-SIGN can get coupled to a similar actomyosin flow in the phagocytic synapse, leading to its early recruitment at (Figure 7).

**Figure 7.**
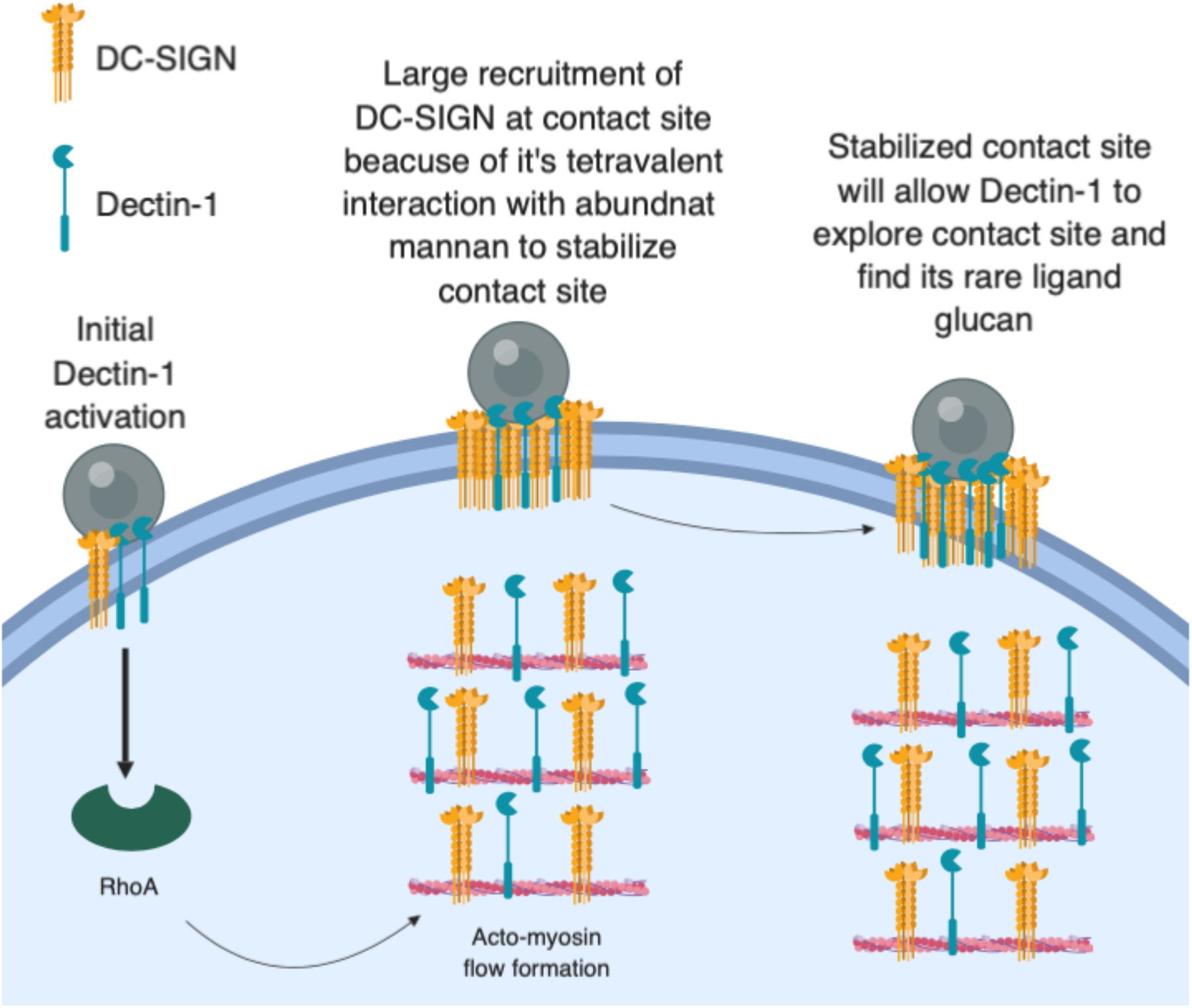
Model for early recruitment of DC-SIGN at fungal contact site. Initial Dectin-1 activation leads to RHOA mediated actomyosin flow formation. DC-SIGN gets coupled to this AMF and start accumulating within contact site, leading to stable capture of fungal particles. After this early stabilization of contact site, Dectin-1 will get time to look for its rare ligand to amplify its signaling, ultimately leading phagocytosis. (Figure created using Biorender)

In contrast to our model, Liu et al. showed constitutive role of microtubule based retrograde transport of DC-SIGN nanoclusters to bring pathogens to the perinuclear region for antigen processing[21]. In this study, DC-SIGN nanoclusters were unladen with pathogen or attached to viral particles. It is possible that microtubule associated transport is a constitutive retrograde transport process involved in receptor recycling or antigen acquisition, but the conditions in Liu et al. did not involve AMF generation because Dectin-1/RHOA axis signaling was absent. Nevertheless, future studies could examine potential the contribution of microtubule mediated transport of DC-SIGN to nascent fungal contacts. Cambi et al. discuss the possible role of DC-SIGN in direct phagocytosis of *C. albicans* by immature dendritic cells [4]. They show enrichment of DC-SIGN within phagosome. This finding is consistent with DC-SIGN’s prominent role in capturing fungal particles, though it also raises possible pro-phagocytic role of DC-SIGN. However, Rosa et al. showed that DC-SIGN plays a role in binding of zymosan particles but is uninvolved in coordinating phagocytic signaling of those particles. Consistent with our findings, Rosa et al. also highlighted the prominent co-localization of actin and Dectin-1 within zymosan contact site, as would be expected for Dectin-1 mediated actomyosin reorganization at the contact site[19].

Geijtenbeek et al. and van Gisbergen et al. showed that DC-SIGN plays a role in intercellular adhesion of DCs with T cells [27,28]. Using an assay of rapid cellular capture of yeast under fluid shear conditions, we found that when Dectin-1 is co-expressed with DC-SIGN, cells could capture significantly more fungal particles than when any of these receptors are expressed individually. Thus, we concluded that the interplay between Dectin-1 and DC-SIGN is important for optimal fungal capture and retention in early fungal phagocytic contacts. ALS5 is a fungal amyloid mannoprotein adhesin which undergoes reorganization into nanodomains under shear. This reorganization of ALS5 under shear exposes binding sites for DC-SIGN, thus making fungal particle sticky for DC-SIGN binding. Thus, it is possible that early, large recruitment of DC-SIGN will improve avidity of interaction with *Candida* through DC-SIGN-ALS5 interactions (or other mannoprotein adhesins) under flow conditions [29], which could be pursued in future research.

In conclusion, we showed that Dectin-1 mediated activation of RHOA-ROCK-Myosin II axis plays important role in active recruitment of DC-SIGN to the *C. albicans* contact site. This is important for capture of fungal particles and the formation of stable host-pathogen contact sites.

## Acknowledgement

We gratefully acknowledge technical advice by Akram Etemadi Amin, Carmen Martinez, Eduardo Anaya and Matthew Graus. The authors declare that they have no conflicts of interest relevant to this work. This research was supported by the University of New Mexico Center for Spatiotemporal Modeling of Cell Signaling (STMC; NIH P50GM085273, AKN) and R01AI116894 (AKN).

## 5. Supplementary data

**Figure S1.**
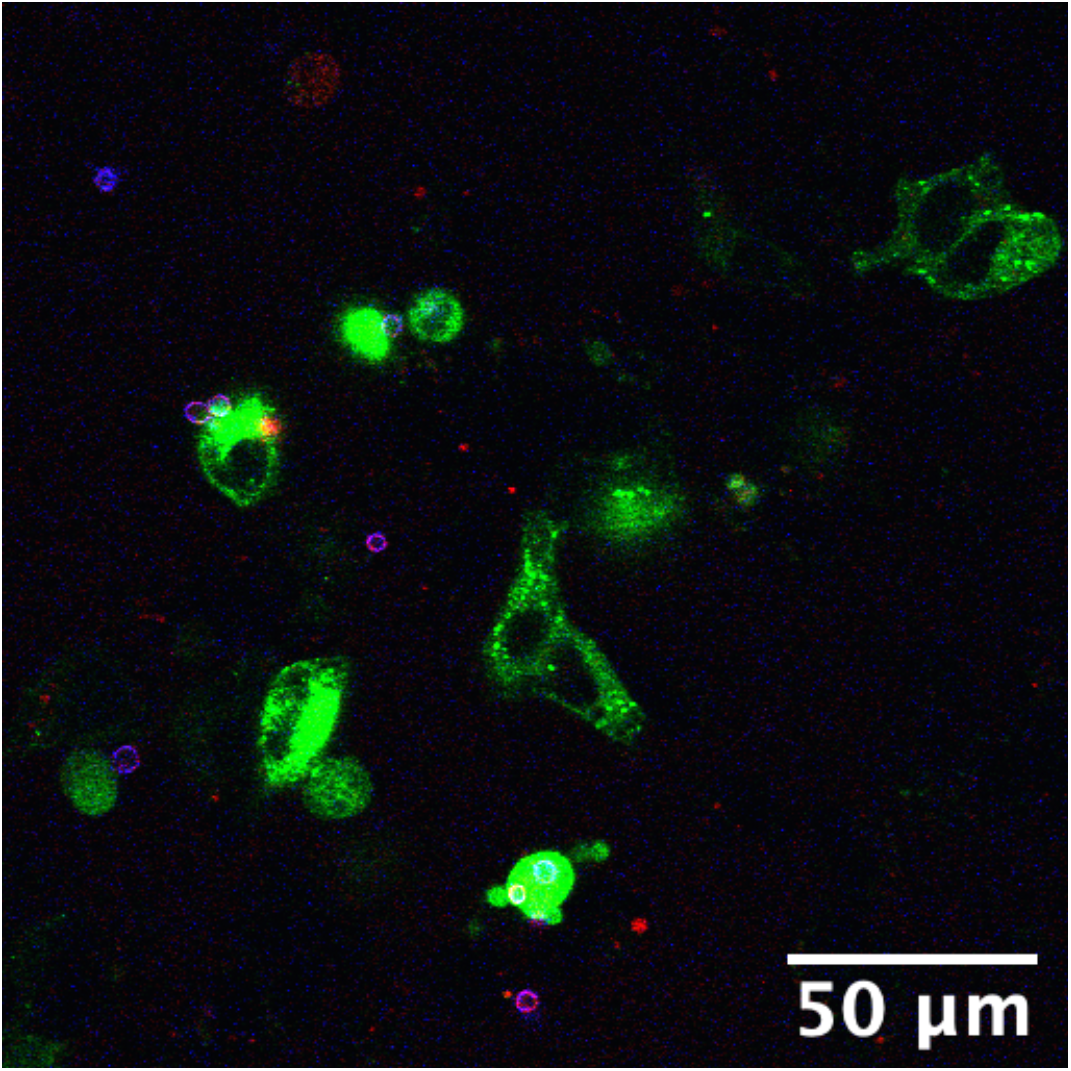
Phagocytosis assay in HEK-293 cells expressing EGFP-DC-SIGN (green) with TRL035 *C. albicans* (blue, calcofluor, cell wall stain; red, cypher5e, pH sensitive phagocytosis indicator). There was no uptake of fungal particles in the DC-SIGN only condition.

**Figure S2.**
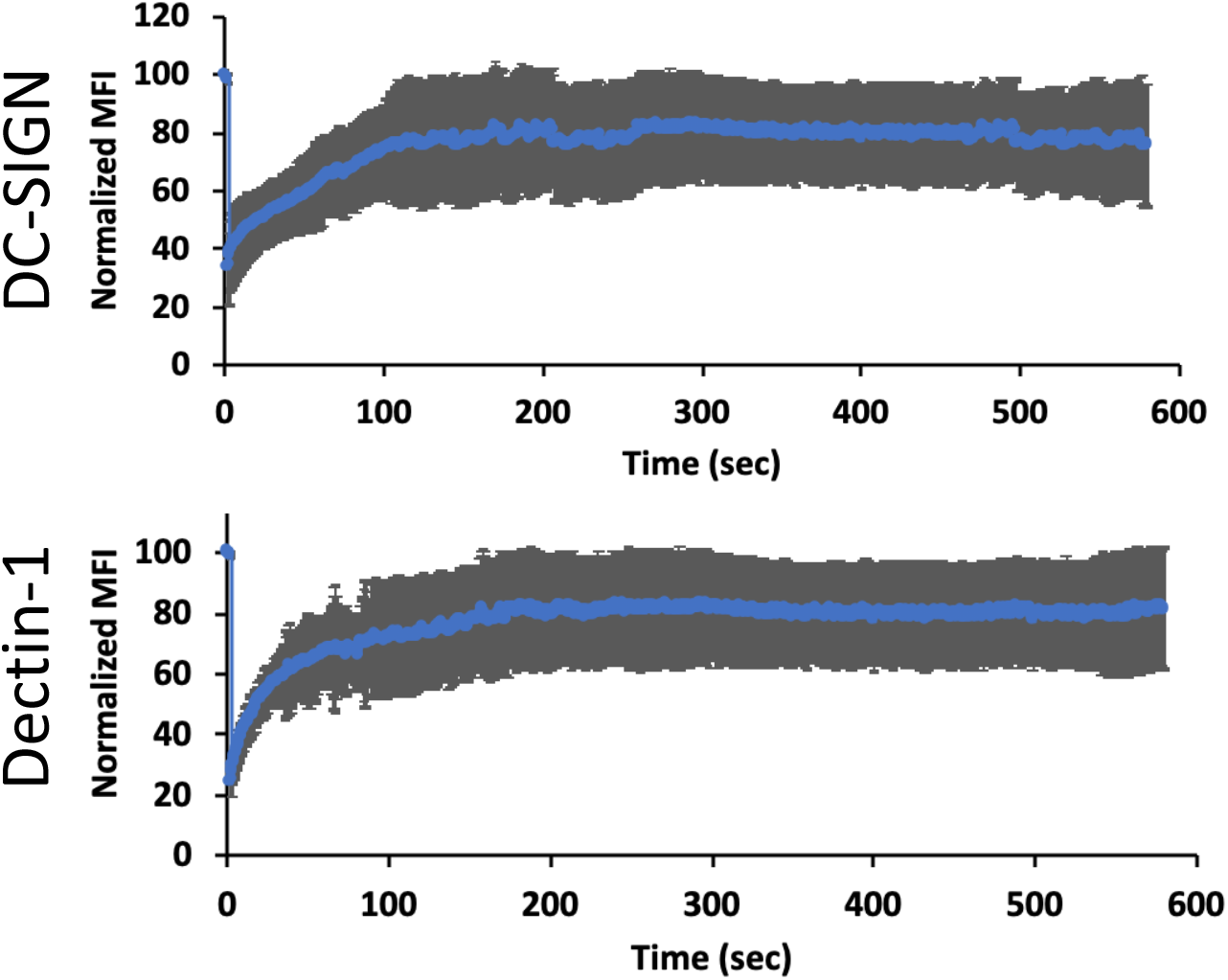
Effect of RHOA inhibitor on DC-SIGN and Dectin-1 mobility in non-contact membranes. (top) DC-SIGN showed 75.72% mobile fraction and recovery t_1/2_ of 35.47 sec. (bottom) Dectin-1 showed 81.77% mobile fraction and recovery t_1/2_ of 14.42 sec

**Figure S3.**
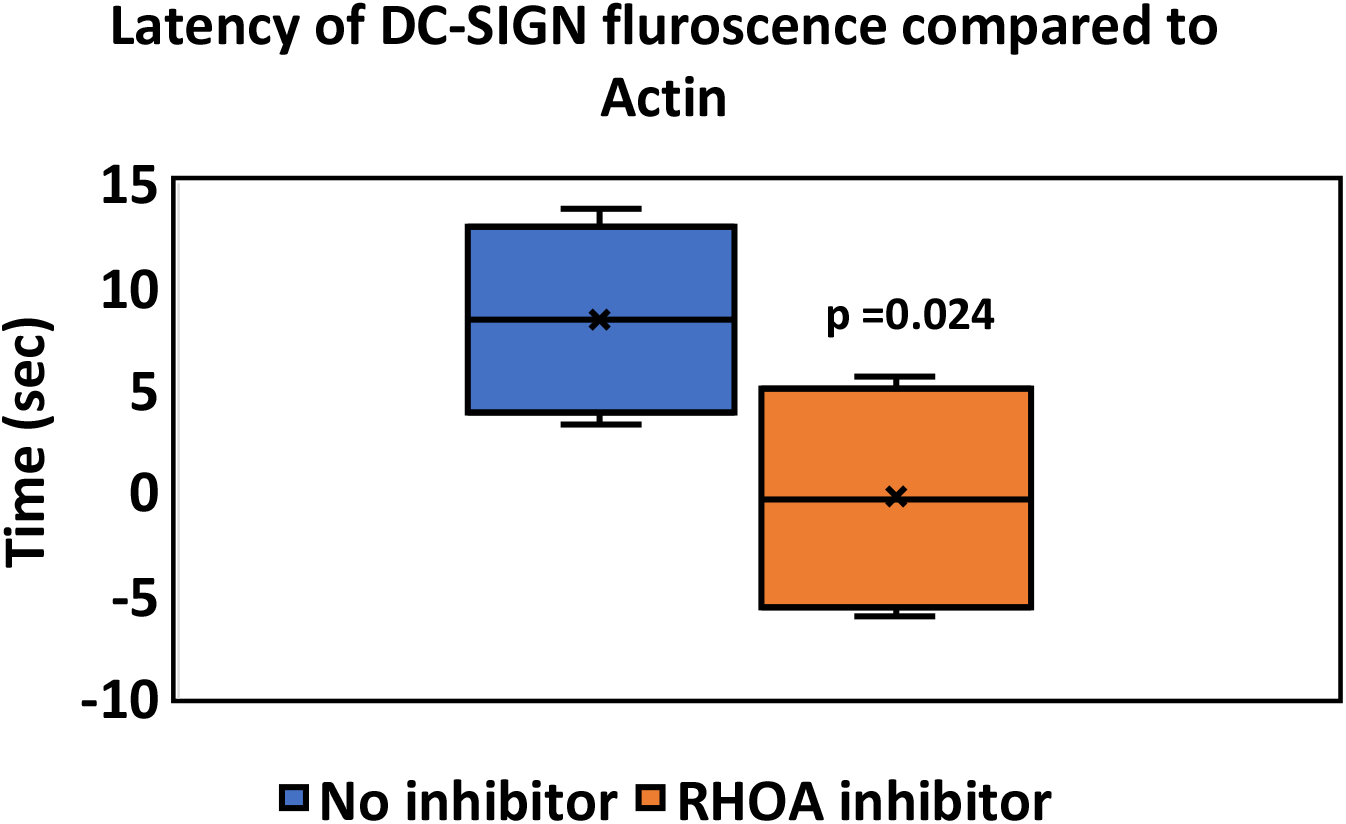
Latency for DC-SIGN compared to F-actin in nascent contact site membranes for no inhibitor and RHOA inhibitor conditions. Box and whisker plot depicts mean (“x”), median (horizontal bar within box), 1^st^ and 3^rd^ quartile (top and bottom of colored box) and minimum and maximum value (whiskers).

### Agent-based modelling: Detailed Methods

For modeling transport of DC-SIGN to fungal contact sites we used Netlogo version 6.1.1. The goal of the simulation was to look at the effects of various modes of lateral transport of DC-SIGN on the kinetics of DC-SIGN recruitment at contact site. By arbitrary convention, we kept world square with the dimension of 5 μm x 5 μm. All other parameters were derived from previously available literature or systematically varied. The essential script for the model is provide below.

#### Model Parameterization

Parameters which were kept constant:

- Contact site diameter was set at 1 μm based on a measured value reported previously[9].
- 1.7 x 10^−4^/nm^2^ mannan binding sites were distributed randomly throughout contact site, based on an estimation of a prominent *C. albicans* mannoprotein previous ly reported [29].
- 2 DC-SIGN tetramers per domain were randomly distributed at domain density of 1.2 x 10^−6^/nm^2^ throughout world area based on prior optical nanoscopy measurements [17,18,23].
- Size of DC-SIGN nanodomain in resting membrane was approximated to 75 nm based on prior optical nanoscopy measurements [17].

Parameters which were varied in model:

- For association and dissociation probability, we ran a separate simulation with single DC-SIGN CRD domains with single mannan moieties available for binding in the world. We optimized model for total simulation time, so that quantity of free and bound fraction of receptor and ligand is stabilized at the end of simulation. We could achieve equilibrium binding for all conditions in 1 hour of simulated time. So, we let model run for each combination of association and dissociation probabilities for 1 hour. Then we calculated K_d_ for each condition using the quantity of unbound ligand, unbound receptor and receptor-ligand concentration. Previous literature has reported experimentally calculated K_d_ for DC-SIGN and mannan interaction to be 50 μM[30]. Hence, we used association and dissociation probabilities which gave K_d_ value between 30 to 60 μM for contact site simulation, bracketing this value. K_d_ values in this range are highlighted as red text in Table 2. Actual values used in simulations for data generation are detailed further below. indicate values around experimentally calculated K_d_ of DC-SIGN-Mannan interaction.
- Probability for DC-SIGN carbohydrate recognition domain to couple (coupling coefficient) to actomyosin flow was varied between 0 to 80%. This was from previously available literature for immunologic synapse [10,22].

**Table 2.** K_d_ value obtained after simulation of each condition of association and dissociation probability for 1 hour. Each condition was simulated in triplicate. Red

|  | Dissoc_prob |  |  |  |  |
| --- | --- | --- | --- | --- | --- |
| Assoc_prob | 0.01 | 0.02 | 0.03 | 0.04 | 0.05 |
| 0.3 | 5.22E-05 | 9.08E-05 | 1.52E-04 | 1.82E-04 | 1.52E-04 |
| 0.35 | 2.18E-05 | 4.00E-05 | 1.01E-04 | 1.11E-04 | 1.21E-04 |
| 0.4 | 1.83E-05 | 3.66E-05 | 1.03E-04 | 9.28E-05 | 9.95E-05 |
| 0.45 | 2.39E-05 | 3.19E-05 | 8.57E-05 | 1.03E-04 | 7.55E-05 |
| 0.5 | 1.40E-05 | 3.39E-05 | 6.54E-05 | 1.11E-04 | 1.11E-04 |
| 0.55 | 1.79E-05 | 3.74E-05 | 6.12E-05 | 1.37E-04 | 1.11E-04 |
| 0.6 | 1.52E-05 | 2.31E-05 | 5.01E-05 | 6.03E-05 | 1.06E-04 |
| 0.65 | 1.09E-05 | 2.59E-05 | 4.71E-05 | 4.87E-05 | 1.52E-04 |
| 0.7 | 1.71E-05 | 2.33E-05 | 4.51E-05 | 3.69E-05 | 5.01E-05 |
| Green < 3E-05 |  | Red 3E-05 to 6E-05 |  | Yellow > 6E-05 |  |

#### Mathematical formulations

- Step sigma for DC-SIGN is based on diffusion coefficient value of 6.5 x 10^−2^ μm^2^/sec for DC-SIGN domain reported by Manzo et al.[18] The mean diffusion radius was obtained from r^2^=4Dt (D=6.5 x 10^−2^ μm^2^/sec, t=0.1s time of a clock tick for model) Hence, r= 161 nm. Therefore, we model diffusion as a random normal jump distance with sigma =161 nm. However, Manzo et al. also measured that 10% of trajectories are immobile, so we modeled this as DC-SIGN domain movements with a 90% probability[18].
- For directed motion component, actomyosin flow velocity of 0.10 μm/sec was derived from previously reported values for immune synapse[10,31].
- With coupling coefficient (*x*), DC-SIGN domains were modeled to be in directed motion of 0.10 μm/sec, fixed heading towards to center of contact site with *x*% probability each time step.
- When not in directed motion, DC-SIGN domains were in random diffusion with random heading and 161 nm diffusion radius in each time step.

#### Running model

- To keep number of DC-SIGNs available in world outside constant, we did allow wrapping of world. Also, to compensate for DC-SIGN which were bound and inside contact site, we replenished the equivalent number of DC-SIGNs to the outside world. This was necessary it would have been computationally infeasible to model the whole cell’s membrane, so we just simulated a small membrane patch. This patch (within the model and outside the contact site zone) is assumed to be freely connected by diffusion to the rest of the cell, which provides an effectively infinite pool of receptor in diffusive equilibrium with the modeled membrane patch. So, we assumed that the concentration of receptor outside contact area should always approximate the entire cell. Even if receptor is lost to the contact site membrane, this would have a negligible impact on the average concentration of receptor in membrane outside the simulation area.
- DC-SIGN immobilization was probabilistic as per association and dissociation probability. So, if DC-SIGN is within binding radius of unbound mannan binding site then probability that it will bind to mannan binding site was as per association probability. Also, bound DC-SIGN dissociated from mannan binding site as per dissociation probability.
- Model itself was run for 10 min. for each condition. This is to keep simulation conditions same as actual experimental conditions.

### Result of contact site simulation

We used only those association and dissociation probabilities which gave K_d_ around 50 μM as indicated by red in table 3. Each simulation replicate consisted of 3 individual simulation runs (at a given parameter set) and the single simulation replicate’s result was considered as the mean behavior of those individual runs. Table 3 reports the mean of these triplicates’ results.

**Table 3.**
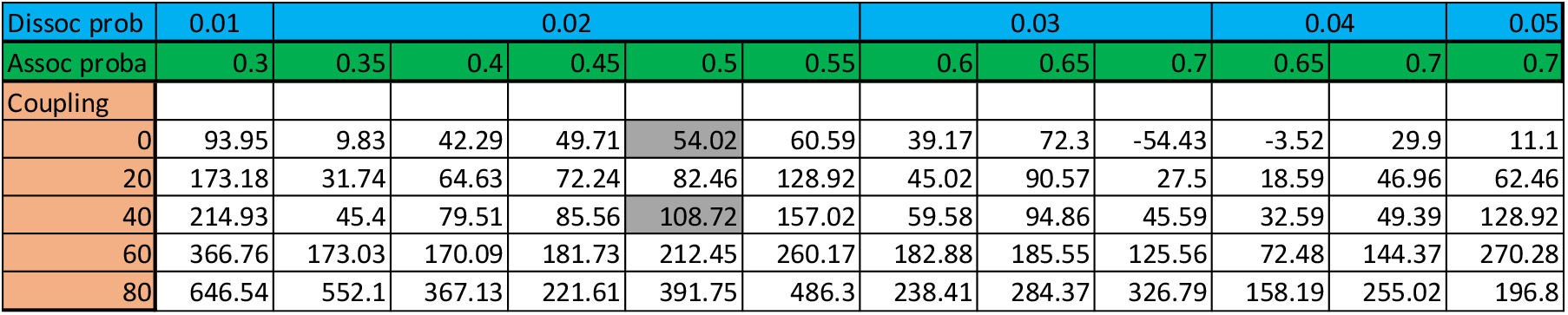
Final increase in DC-SIGN densities compared to pre-contact densities at the end of 10 min. simulation for each specific condition of association probability, dissociation probability and coupling coefficient. Each condition was simulated in triplicate.

We found that dissociation probability of 0.02 and association probability of 0.5 with coupling of 40% gave increase in DC-SIGN density of 108.72% (highlighted in gray in table 3.3), this is similar to the experimental value of 103.15% obtained with DC-SIGN and Dectin-1 co-expression contact sites (Fig. 3.1e).

This same condition of dissociation probability of 0.02 and association probability of 0.5 with 0 coupling gave increase in DC-SIGN density of 54.02% (highlighted in gray in table 3.3). This is similar to what we obtained with RHOA, ROCK and Myosin-II inhibitors which showed DC-SIGN recruitment of 41.47%, 62.96% and 44.04% respectively (Fig. 3.3).

Overall, in table only dissociation probability of 0.02 and association probability of 0.5 gave a global best fit to what we observed experimentally.

### Model Script: Main Simulation

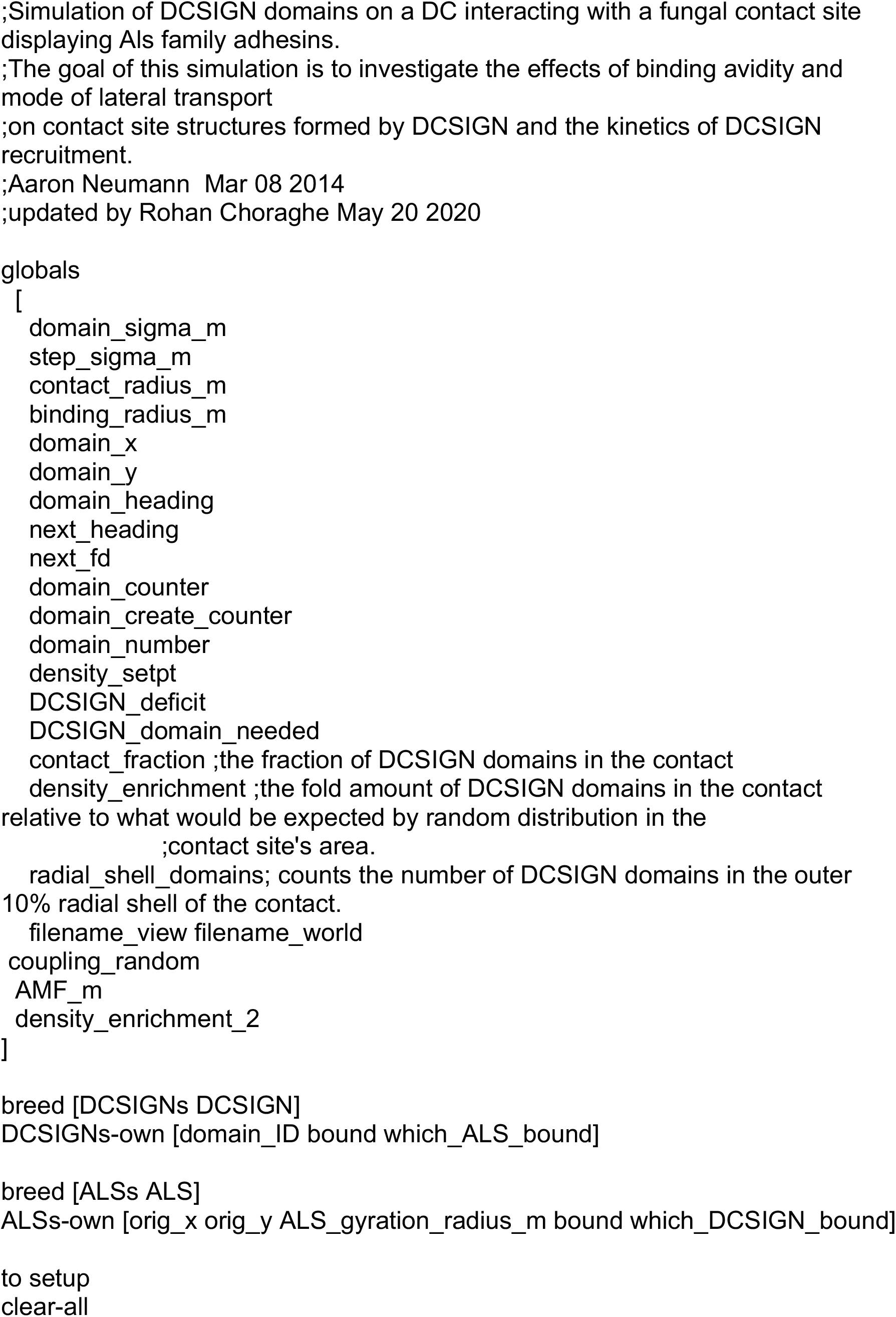

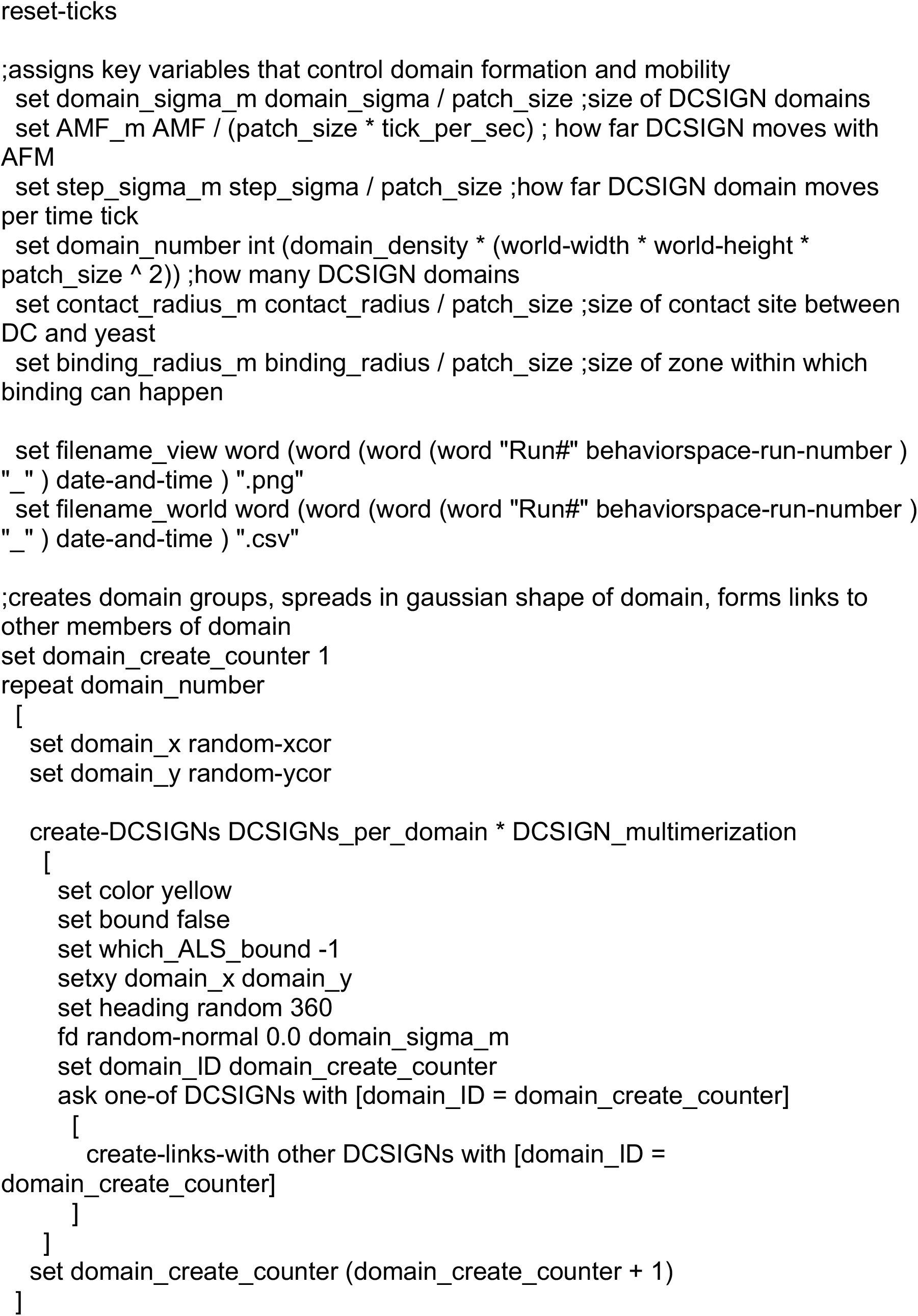

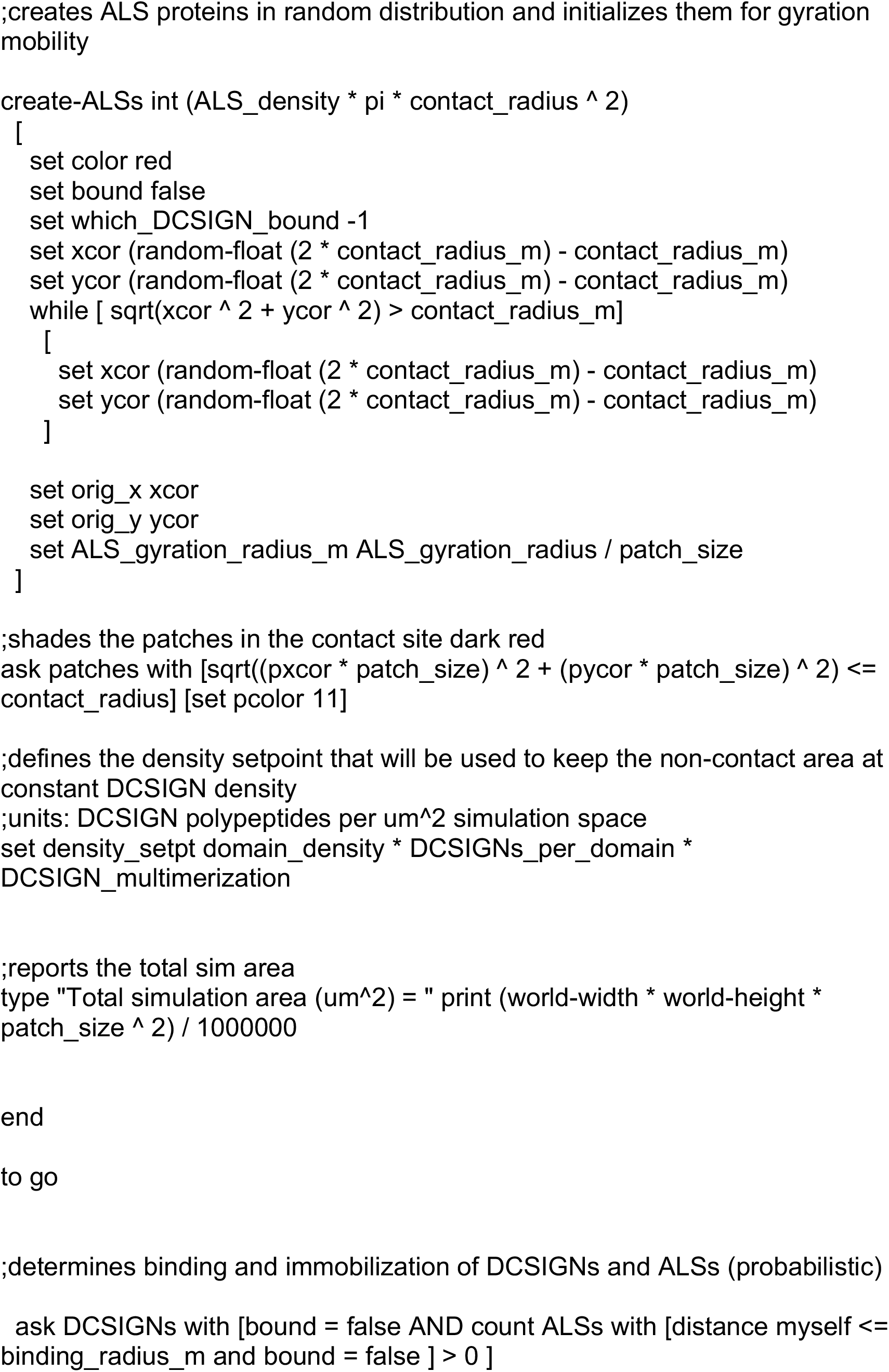

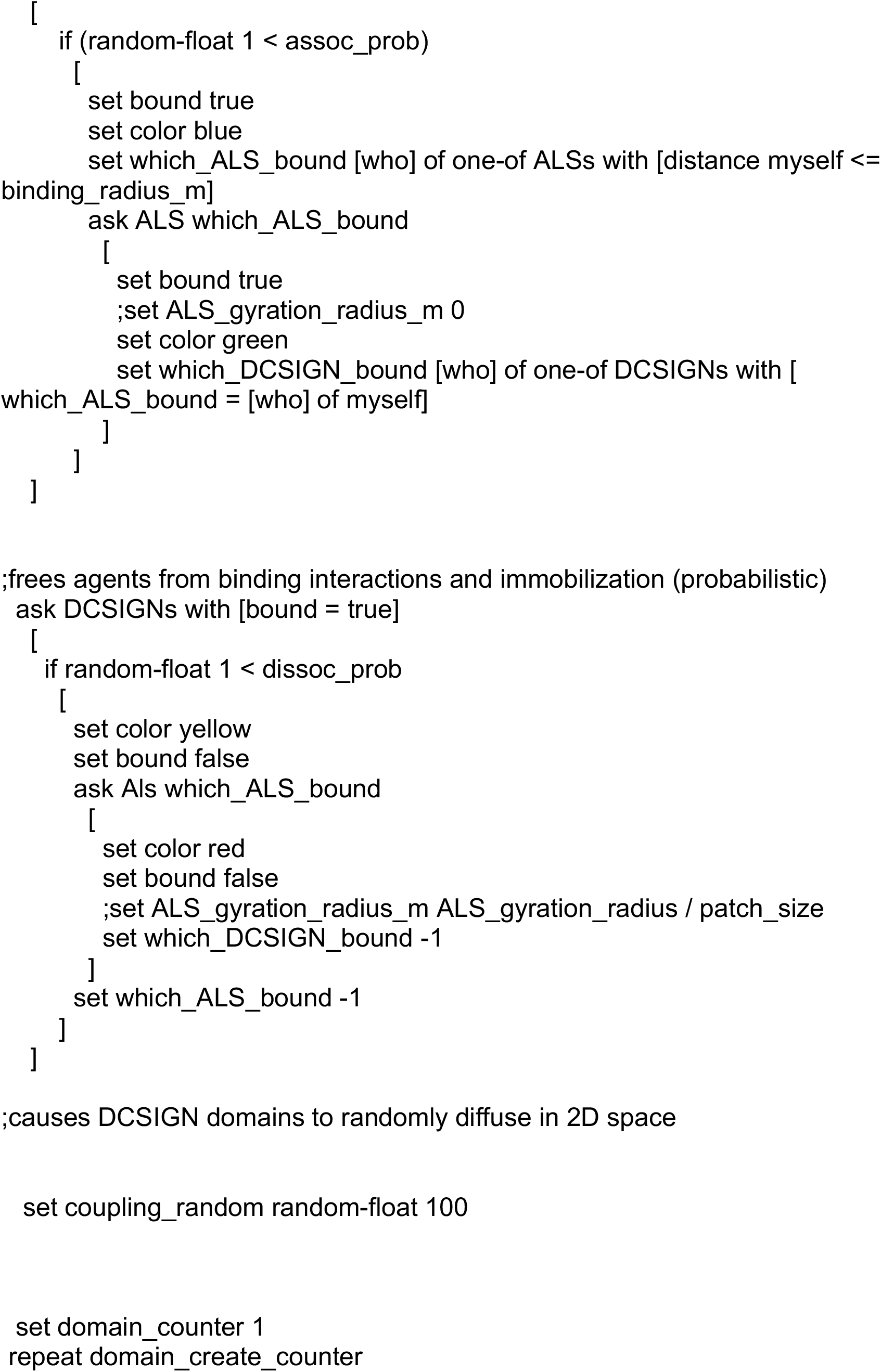

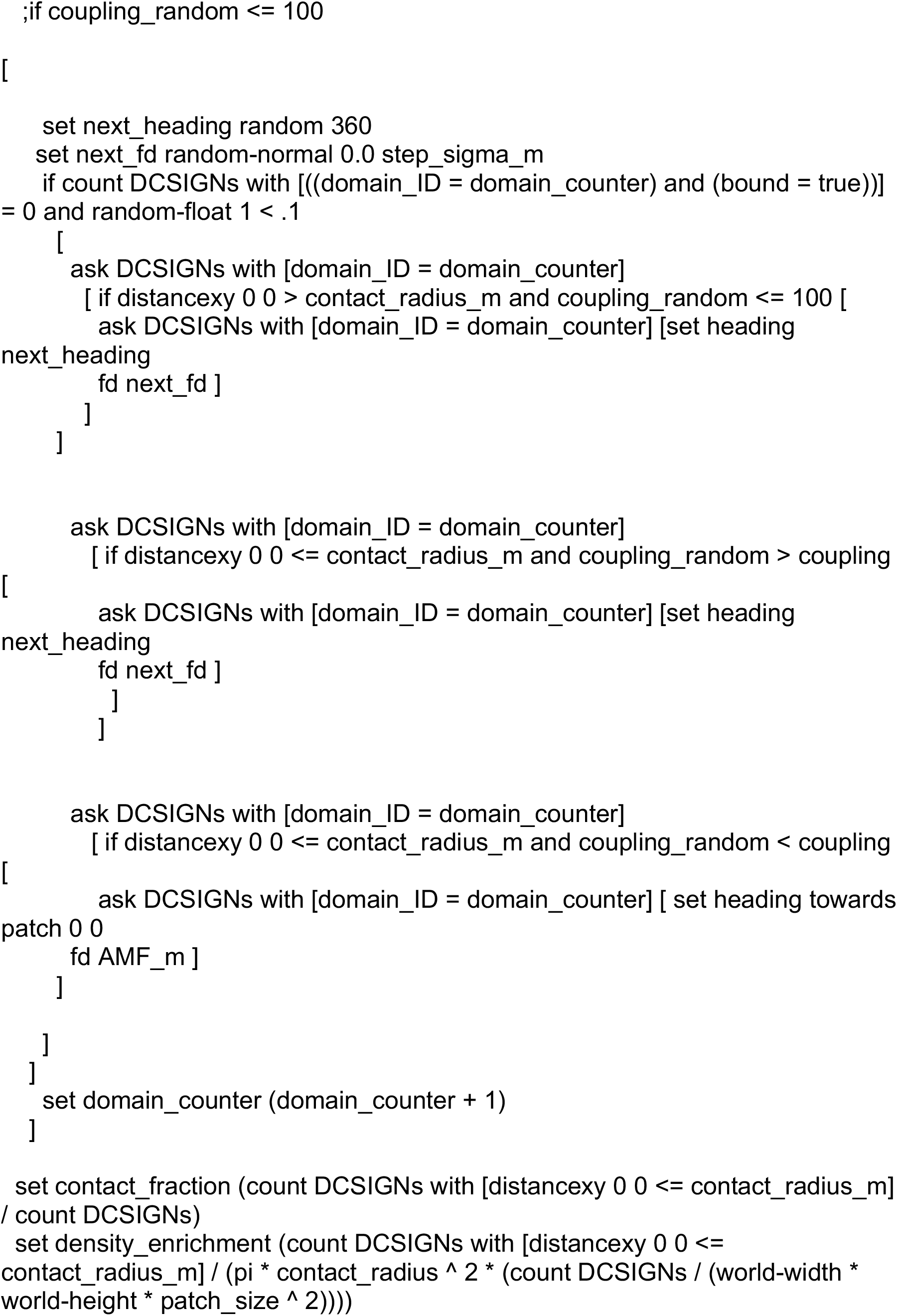

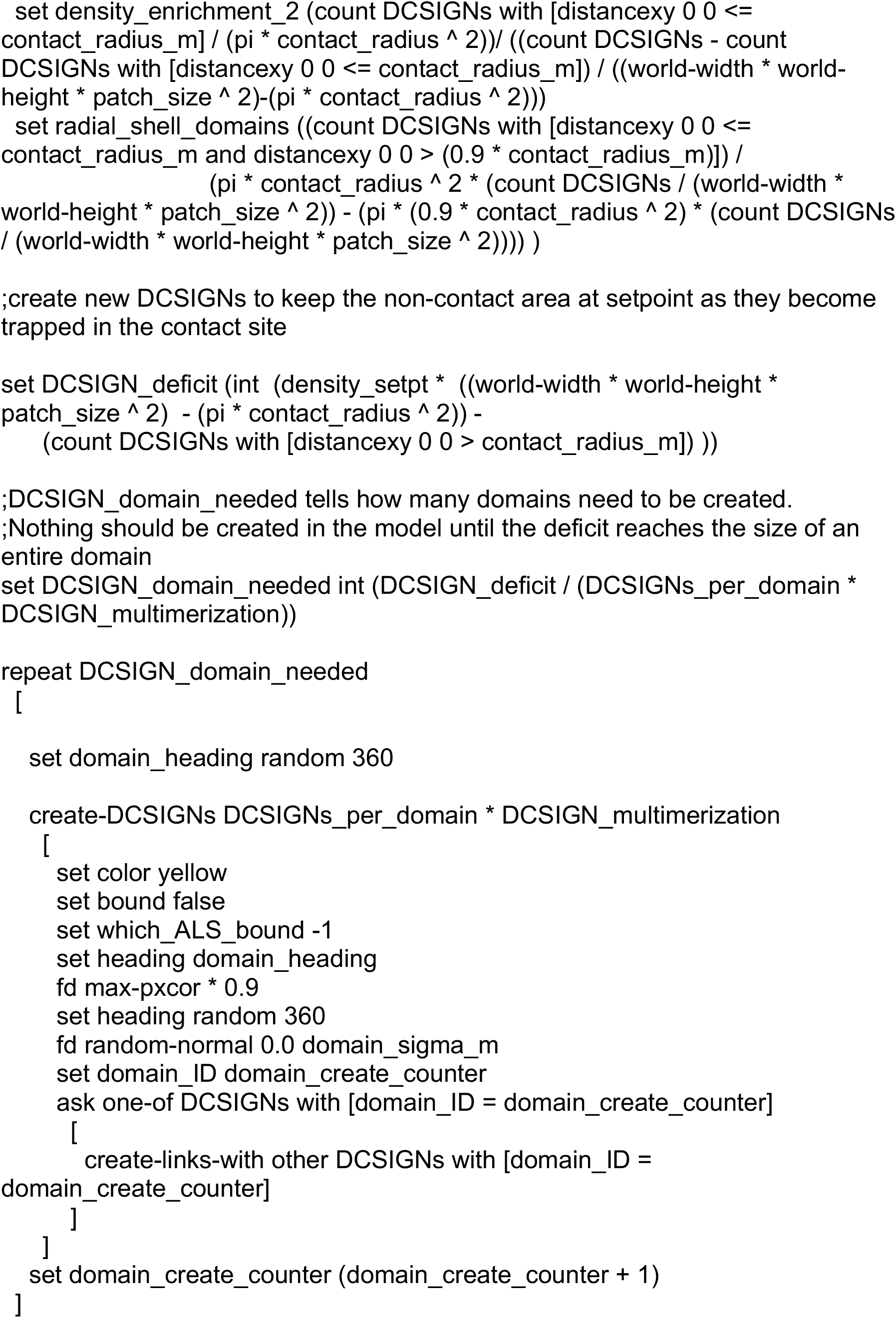

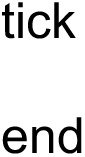

### Model Script: Kd estimation

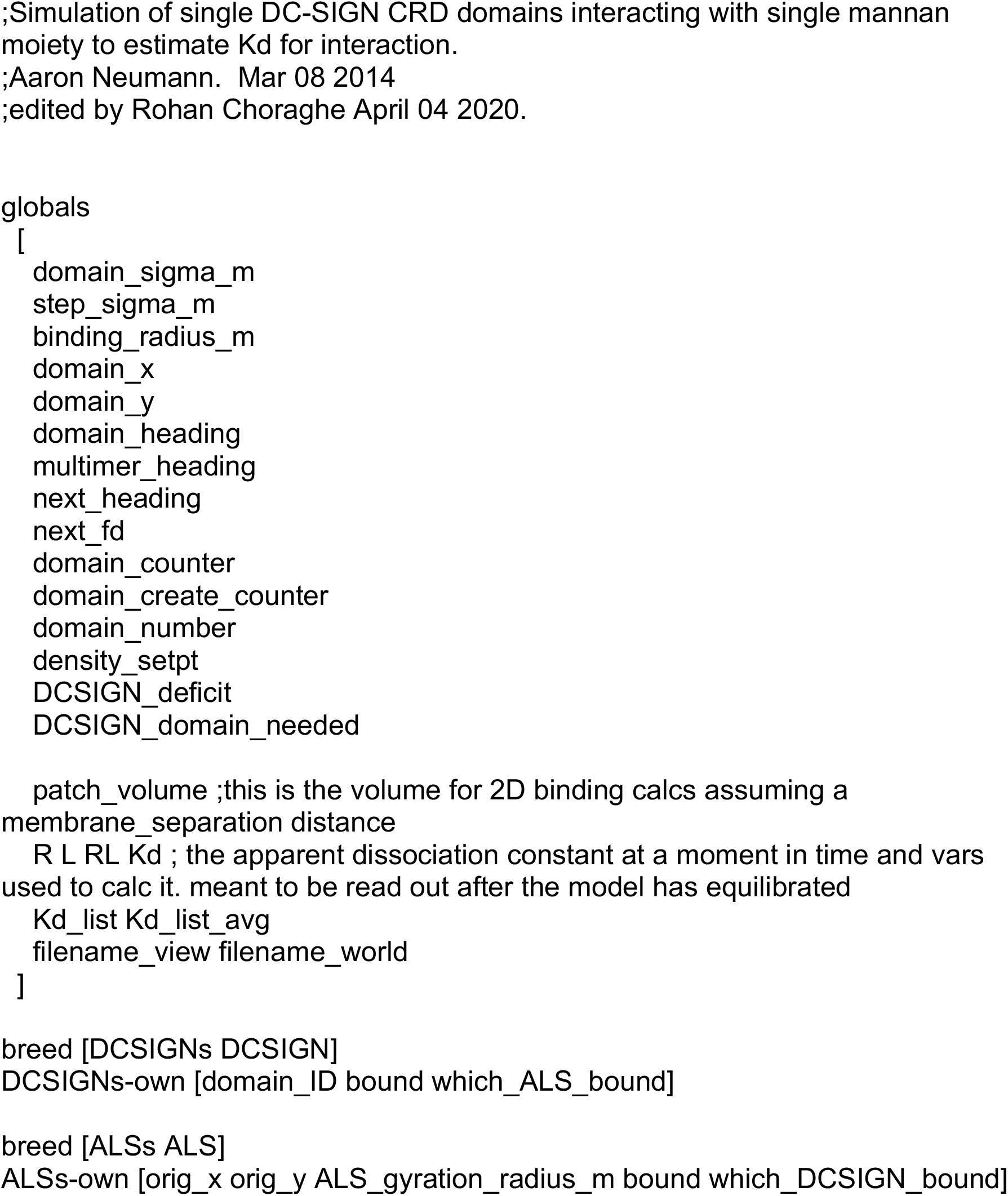

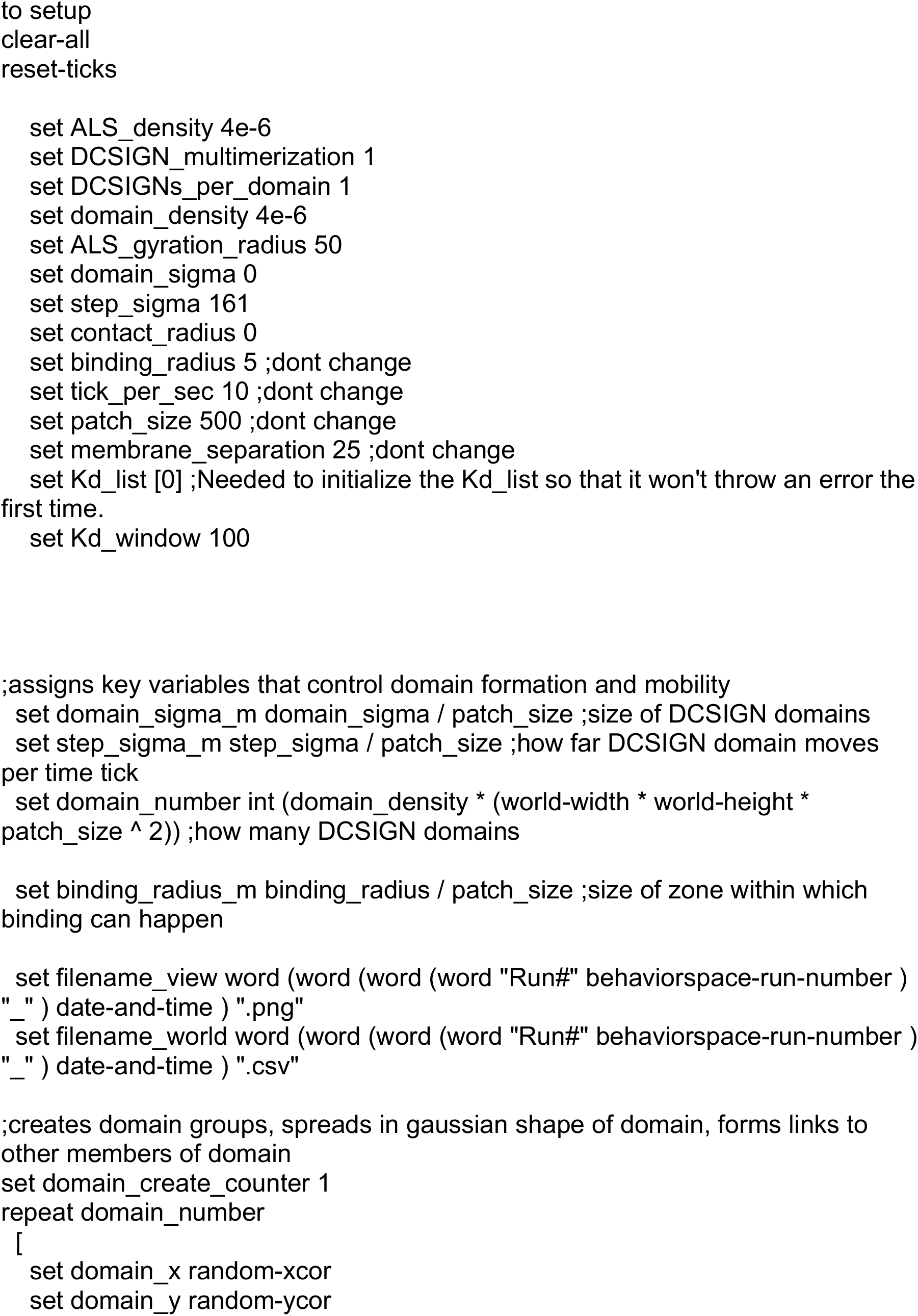

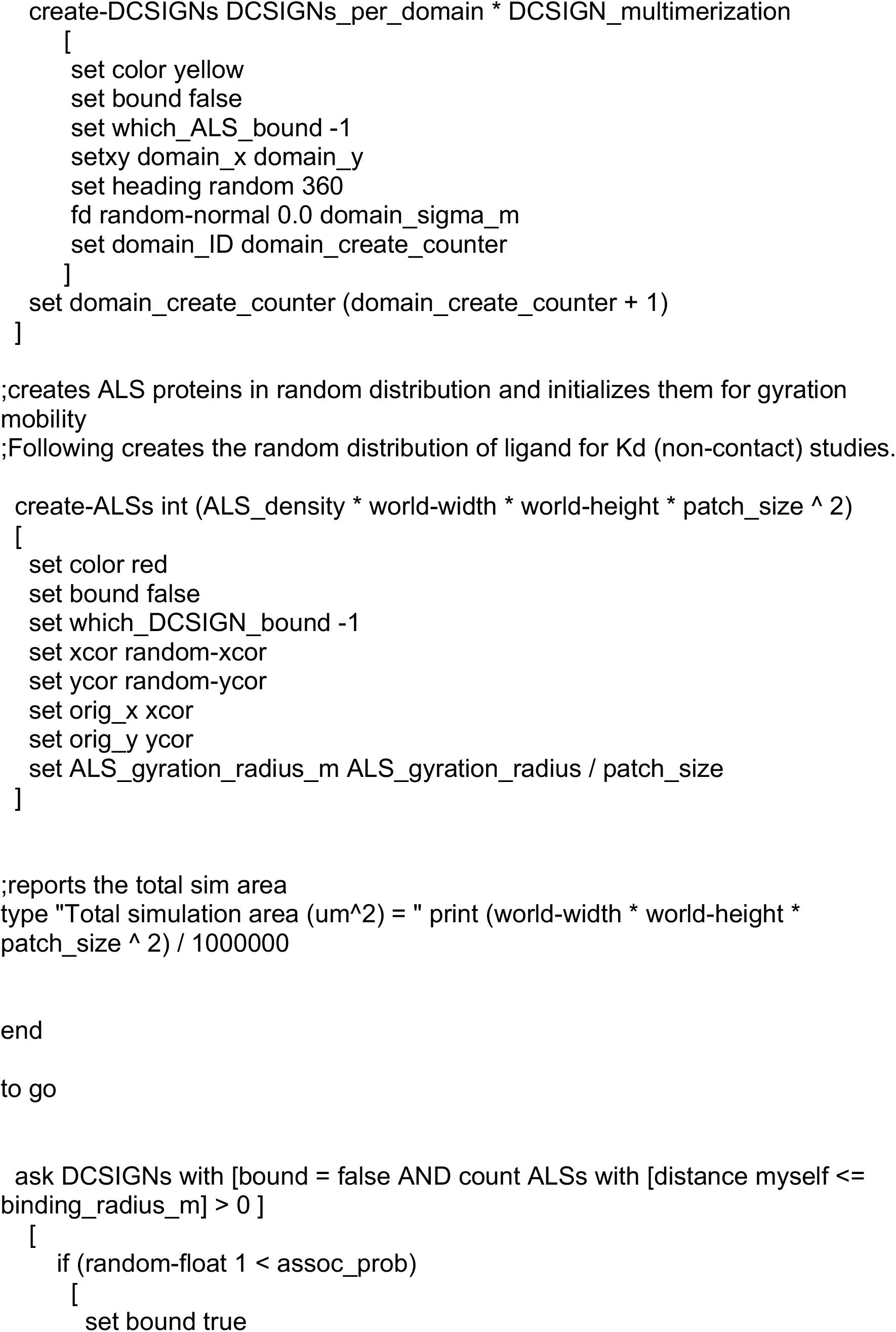

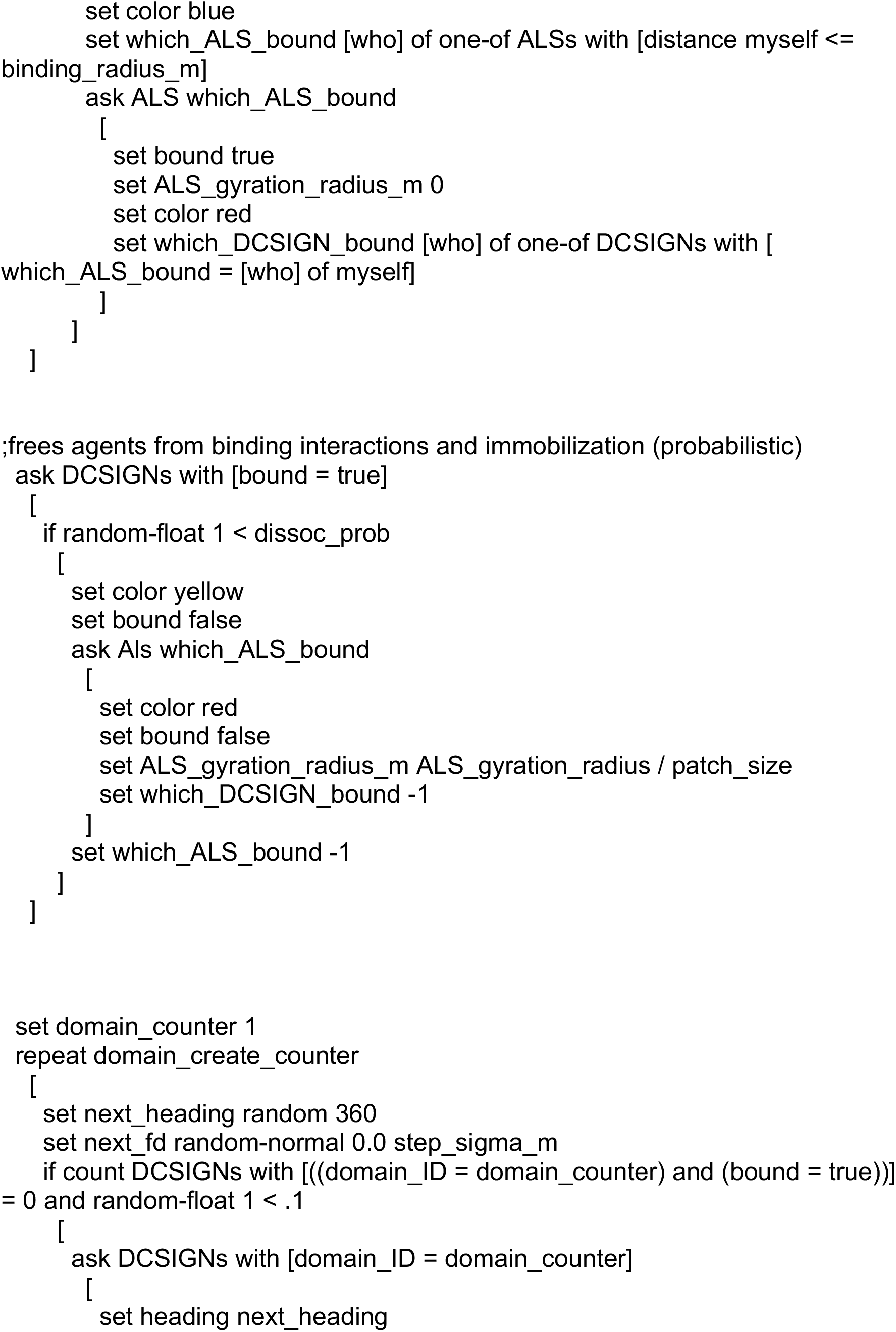

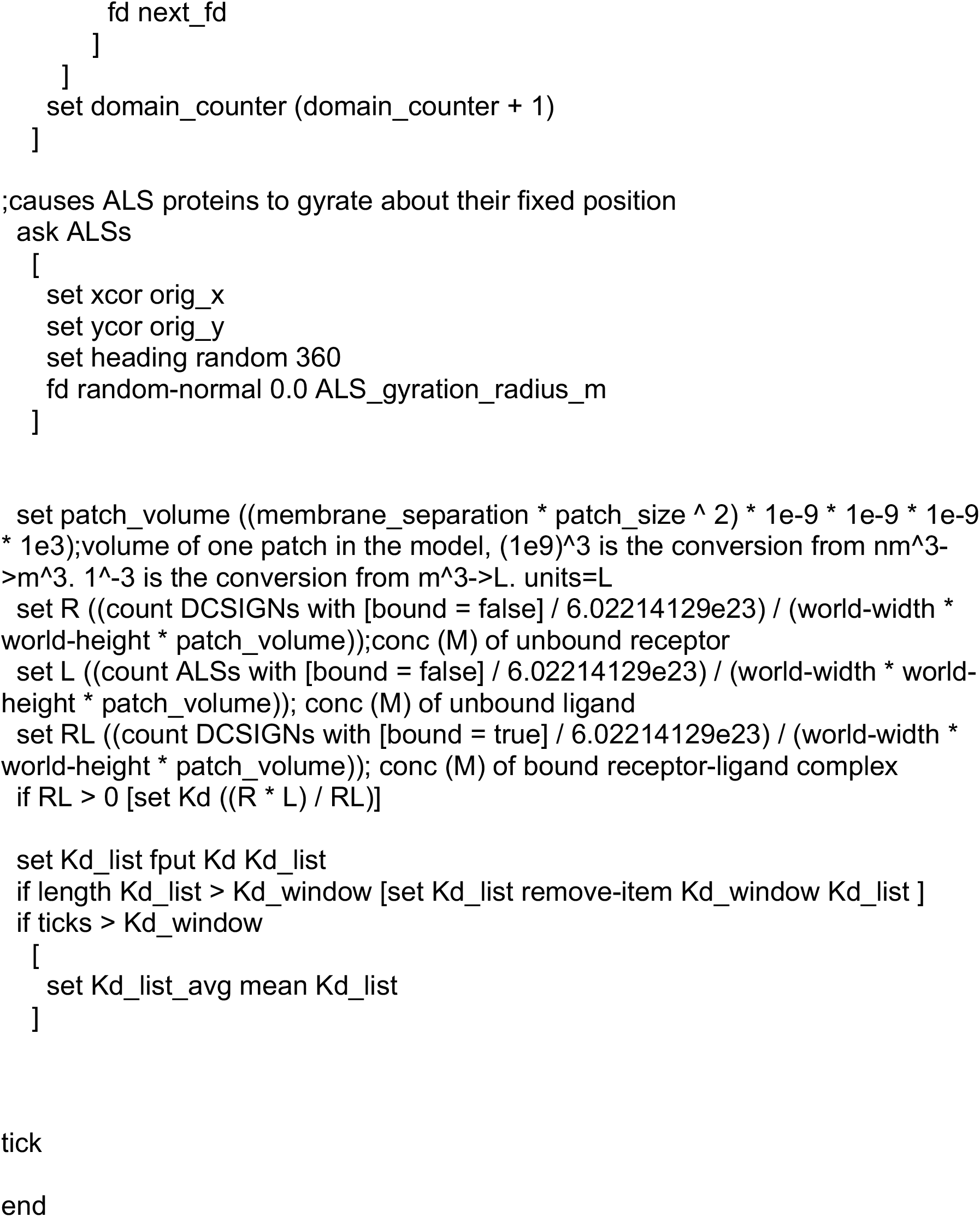

## Notes

### Competing Interest Statement

The authors have declared no competing interest.

